# Germline activity of the heat shock factor HSF-1 programs the insulin-receptor *daf-2* in *C. elegans*

**DOI:** 10.1101/2021.02.22.432344

**Authors:** Srijit Das, Sehee Min, Veena Prahlad

## Abstract

The mechanisms by which maternal stress alters offspring phenotypes remain poorly understood. Here we report that the heat shock transcription factor HSF-1, activated in the *C. elegans* maternal germline upon stress, epigenetically programs the insulin-like receptor *daf-2* by increasing repressive H3K9me2 levels throughout the *daf-2* gene. This increase occurs by the recruitment of the *C. elegans* SETDB1 homolog MET-2 by HSF-1. Increased H3K9me2 levels at *daf-2* persist in offspring to downregulate *daf-2,* activate the *C. elegans* FOXO ortholog DAF-16 and enhance offspring stress resilience. Thus, HSF-1 activity in the mother promotes the early life programming of the insulin/IGF-1 signaling (IIS) pathway and determines the strategy of stress resilience in progeny.

**One Sentence Summary:** HSF-1 recruits MET-2 to silence *daf-2* and mediate early life programming of *C. elegans* upon stress

---

Many organisms display transgenerational plasticity whereby the memory of parental exposure to environmental stress is retained through meiosis and cycles of mitosis to alter later life cellular responses and survival mechanisms of offspring (*1–4*). This early life programming, or maternal effect, can be beneficial for progeny, altering their phenotypes to decrease predation risk (*5*); alternatively, it can be detrimental as has been observed in humans where maternal stress can increase the late-life suceptibility of offspring to metabolic and neuropsychiatric disease (*6–9*). In the free-living nematode *Caenorhabditis elegans* too, epigenetic mechanisms that affect chromatin in one generation alter offspring lifespan and behavior in subsequent generations (*10–30*). However, in all cases, how epigenetic changes arise during perinatal life is not clear.

As in many organisms, in *C. elegans*, environmental stressors that impact the parent also affect the unfertilized gametes. In the *C. elegans* parent, stress activates the heat shock transcription factor, HSF1 (HSF-1 in *C. elegans*), which increases the expression of protective heat shock protein (*hsp*) genes to counteract stress-induced damage (*31, 32*). We had previously shown that even a brief and transient exposure to increased temperatures activates HSF-1 in the germline of *C. elegans*, and that this rapid activation of HSF-1 is essential to maximize the animal’s reproductive output (*33*). Here we investigated whether HSF-1 activity in the maternal germline of *C. elegans* contributed to early life programming of offspring to alter their later-life strategies of survival.

To do this, we first established an experimental protocol to activate HSF-1 in germ cells of *C. elegans* and examine the later-life phenotypes of offspring generated from these germ cells (Fig. S1A). We exposed one-day-old mothers to 34°C heat-shock for varying durations. During this stress exposure, oocytes resident in the animal’s germline as part of the germline syncytium also experienced heat-shock. We then allowed these oocytes to complete maturation, undergo fertilization, and develop into adults at normal growth temperature of 20°C. To activate HSF-1 in the germline, we chose regimen of long heat exposure (30 minutes or 60 minutes to 34°C) that is typically used to activate HSF-1 throughout the animal (*33–36*) and a shorter, 5-minute heat-shock at 34°C, which we have previously shown is sufficient to induce HSF-1 transcriptional activity (*33*). By determining the number of fertilized eggs in the uterus and unfertilized oocytes in the germline (8.6 ± 0.5/animal) after stress exposure, and egg laying rates of the mothers (Fig. S1 B, C), we established that embryos collected 2 hours after a 5 minute maternal heat shock, 4 hours after a 30 minute heat shock, or 8 hours after a 60 minute heat-shock, were embryos that were reliably generated by oocytes in diakinesis that were resident in the maternal germline during heat exposure, but fertilized at 20°C during maternal recovery (Fig. S1 B, C). Therefore, embryos laid 2-4 hours, 4-8 hours or 8-12 hours following a 5 minute, 30 minute or 60 minute maternal heat shock respectively, were used for all experiments (Fig. S1A). These embryos were allowed to develop into one-day-old adults at 20°C, subjected to a severe heat shock at 37°C, and scored for survival to determine whether early life stress exposure had altered their later life stress resilience. We found that compared to progeny that developed from oocytes that had not been subjected to heat stress, progeny that developed from oocytes that had been exposed to heat-shock while in the maternal germline were more resilient to the subsequent exposure to severe heat stress and significantly more survived (Fig. 1A). Moreover, while embryos collected between 2-4 hours post-5 minute maternal heat shock were more stress resistant, oocytes fertilized and laid later after maternal stress (collected 8-12 hours post 5 minute maternal heat shock) did not display this heightened thermotolerance (Fig. 1A), suggesting that the early life programming of stress resilience was occuring through oocytes, and not sperm, since the same heat-shocked sperm also fertilize these later-laid embryos.

**Figure 1:**
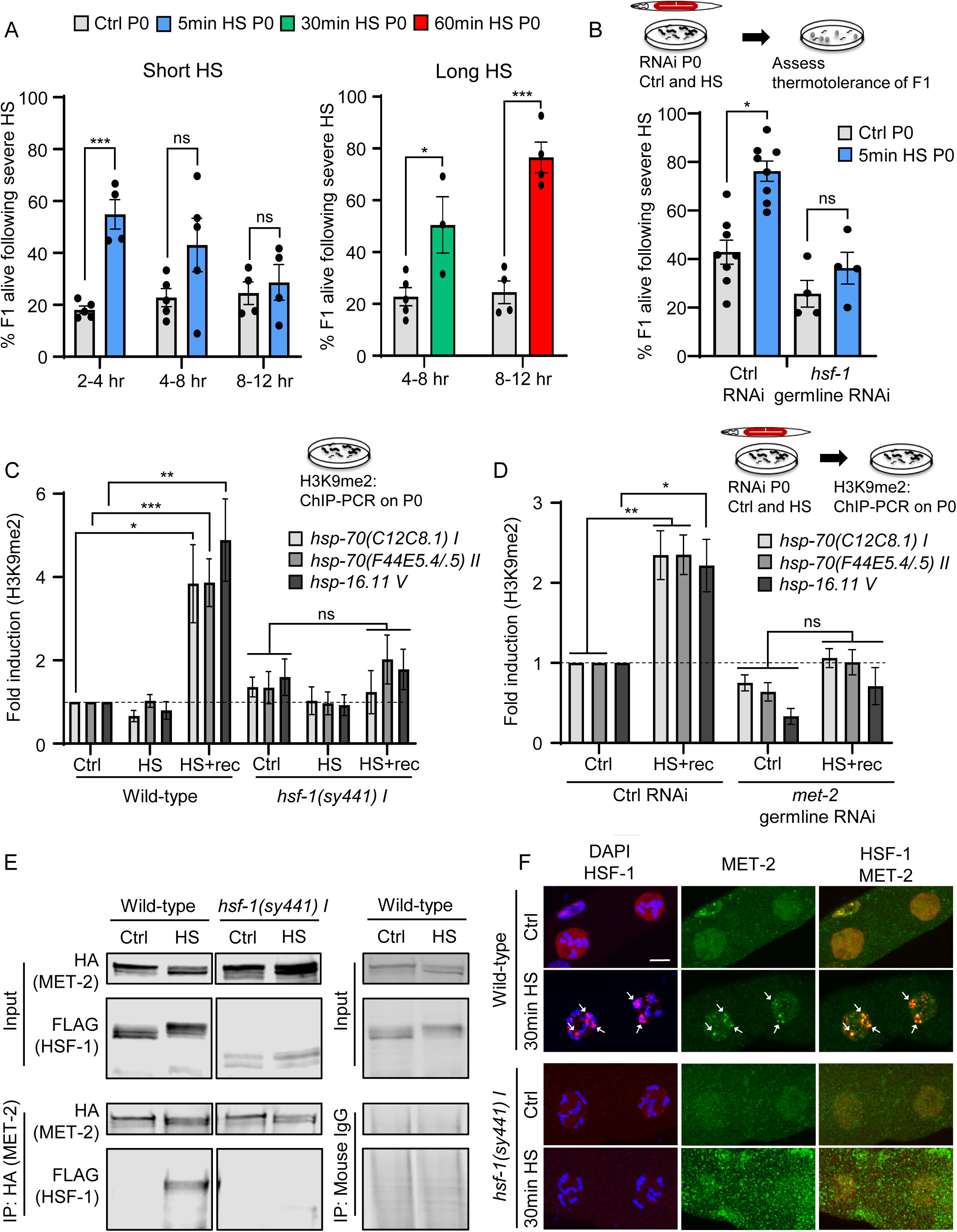
Germline activity of HSF-1 promotes the recruitment of MET-2 to HSF-1 target genes and alters the stress resilience of progeny. **A,** Stress resilience of progeny (F1) of day one *C. elegans* mothers subjected to a short (5 minute) or long (30 minute or 60 minute) heat shock at 34°C. X-axis: the time interval at 20°C <u>after</u> the maternal heat-shock at which progeny were collected. Y axis: Percent F1 progeny that surive severe heat-stress. Legend indicates the duration of maternal heat shock. All the progeny laid by control and heat-shocked mothers within the given time frame were assessed for survival after severe heat stress. n=3-6 experiments/ time point. Each experiment represents 5-15 P0 mothers/ condition/time interval and an average of 24.3±2.3 F1 progeny/time interval/experiment for the short heat shock experiments and 50.4±7.8 F1 progeny/time interval/experiment for longer heat shock experiments. **B,** Stress resilience of F1 progeny of control mothers (Ctrl), and mothers subjected to the short (5 minute) heat shock following the RNAi mediated downregulation of *hsf-1* only in the maternal germline. L4440 empty vector was used as the control. n=4-8 experiments. Each experiment represents thermotolerance of all the progeny laid within the 2-4 hours post-heat shock (average F1 progeny from mothers subjected to germline knockdown using control L4440 RNAi and *hsf-1* RNAi: 28.4±1 and 24.1±1.3 /experiment respectively). **C,** H3K9me2 occupancy (relative to that in control wild-type animals) at the promoter proximal 5’-UTR regions of *hsp-70 (C12C8.1)I, hsp-70 (F44E5.4/.5)II* and *hsp-16.11 V* in P0 wild-type animals and *hsf-1 (sy441)I* animals under control conditions (Ctrl), immediately (HS), and 2 hours after (HS+rec) a 5 minute heat shock (n=4-7 experiments). See Figure S2 D and Material and Methods for details regarding the 5’-UTR regions assayed. **D,** H3K9me2 occupancy (relative to control wild-type animals) at the promoter proximal 5’-UTR regions of *hsp-70 (C12C8.1)I, hsp-70 (F44E5.4/.5)II* and *hsp-16.11 V* in P0 animals subjected to RNAi induced kncokdown of *met-2* only in the germline. L4440 empty vector was used as the control. H3K9me2 occupancy was assessed in control animals (Ctrl) and at 2 hours recovery post-5’minute heat shock (HS+rec), corresponding to the time point where H3K9me2 levels increase at these regions (Fig 1C). (n=3-10 experiments). **E,** Control and heat-shocked (30 minutes) day one wild-type and *hsf-1(sy441) I* mutants expressing FLAG::HSF-1 and MET-2::HA were immunoprecipitated with anti-HA antibody. Wild-type animals were also immunoprecipitated using control IgG antibodies. Proteins were separated by SDS/PAGE, and western blot analysis was performed using the antibodies indicated on the left. Representative Western blot shown. 10% of Input is included. (n=3 experiments). **F,** Representative micrographs showing projections of confocal z-sections through nuclei of oocytes in diakinesis in wild-type animals and *hsf-1(sy441)I*, under control conditions and upon heat-shock (30 minutes; also see **Fig. S6**). Animals express HSF-1 tagged at its endogenous locus with an N-terminal 3X FLAG and MET-2 tagged at its endogenous locus at the C-terminus with 3X HA. Arrows indicate HSF-1 and MET-2 colocalization in nSBs at condensed chromosomes (DAPI) upon heat shock. Panels from left to right show: overlap of HSF-1 immunostaining (red) with DAPI (blue), MET-2 immunostaining alone (green), and overlap of HSF-1 (red) and MET-2 (green) immunostaining. Scale bar:5µm. A-D: Data show Mean ± Standard Error of the Mean *, *p*<0.05; **, *p*< 0.01 ***, *p*<0.001; ns, non-significant; (A-D:: ANOVA with Tukey’s correction; Unpaired Student’s t-test).

We confirmed that heat-shock exposure activated HSF-1 in the germline by using ChIP-PCR (Chromatin immunoprecipitation followed by PCR) to measure the occupancy of HSF-1 at the promoter regions of *hsp* genes in chromatin extracted from the whole animal after *hsf-1* was knocked down only in the germline using RNA interference (RNAi; Fig.S2 A-D). For these experiments, animals that expressed endogenous HSF-1 tagged with 3X FLAG were used and HSF-1 occupancy at the *syp-1* promoter (Fig S2 C) served as a control. Germline knockdown of *hsf-1* abolished HSF-1 binding to target *hsp* promoter regions upon a 5-minute heat shock, and HSF-1 binding was still reduced upon longer durations of heat exposure (Fig. S2 A, B). This confirmed that HSF-1 is indeed activated in the germline cells upon heat shock, and moreover during the first 5-minutes of heat exposure, HSF-1 binding to target *hsp* genes occurs almost exclusively in germline cells.

Resilience to heat stress depends on molecular mechanisms that modulate protein homeostasis. Consistent with their ability to better survive severe temperature stress, progeny of mothers exposed to heat stress also partially suppressed the accumulation of misfolded and aggregated polyglutamine (polyQ) expansion proteins that begin to aggregate later in life (Fig. S3 A). This occurred even though steady-state expression levels of polyQ expansion proteins in progeny of stressed mothers were similar to that in offspring of non-stressed mothers (Fig. S3 B). Thus, in addition to increasing the ability of progeny to survive heat-stress, early life stress exposure also altered the cellular mechanisms of protein homeostasis in offspring.

Downregulation of *hsf-1* only in the maternal germline using RNAi was sufficient to abolish the early life programming of stress tolerance in offspring (Fig. 1B) indicating that germline HSF-1 was required for the increased thermotolerance of progeny of stressed mothers. We also tested whether the early life programming of stress tolerance occurred through the stress-induced activity or the basal activity of HSF-1 in the germ cells by measuring the stress resilience of animals harboring a mutation in *hsf-1, hsf-1 (sy441) I* (37). These animals express a truncated HSF-1 protein lacking the transactivation domain, where HSF-1 binds constitutively to the promoter regions of its target genes in the absence of heat shock, but is deficient in heat-shock induced binding (Fig. S3 C, D) and transcription (*33*). Progeny that developed from heat-shocked germ cell oocytes of *hsf-1 (sy441) I* animals also did not undergo early life programming and did not display enhanced stress resilience (Fig. S3 E). These data showed that stress induced activity of HSF-1 in progeny when they were resident as oocytes in the maternal germline was altering their stress resilience even after they had undergone fertilization and developed into adults.

During normal development, chromatin modifications aquired during germline transcriptional activity are removed at fertilization by histone modifying enzymes to epigenetically reprogram the embryo (*4, 14, 38*). In *C. elegans,* this occurs during the passage of germ cells through the maternal germ line by the activities of H3K4 demethylases SPR-5 and RBR-2, which remove H3K4 marks of active gene expression that are added during normal germline transcription (*26–29*). These demethylase enzymes synergize with the histone H3 lysine 9 (H3K9) SET domain methyltransferase MET-2 (*26, 39–41*), responsible for the addition of repressive H3K9me2 modifications and together erase the memory of germline transcription of spermatogenesis genes to prevent their ectopic expression in the next generation (*26–29*). Since adult offspring of stressed mothers were more stress resilient, we first considered the possibility that HSF-1 germline transcription had escaped germline reprogramming, allowing HSF-1 to preserve the memory of prior transcription and more readily induce stress-protective genes in the next generation upon their exposure to stress. To test this, we investigated whether stress-induced *hsp* expression in adults that developed from oocytes subjected to a 5-minute heat-shock while in the maternal germline was enhanced when compared to that in adults from control non heat-shocked mothers. However, this was not the case; instead of being enhanced, heat-shock protein gene expression in adults that developed from heat-shocked oocytes was attenuated. This was evident from the lower levels of *hsp70* mRNA (Fig. S4 A, B), as well as from the decreased binding of HSF-1 to the promoter regions of *hsp* genes (Fig. S4 C, D) in the offspring from heat-shocked mothers, when compared to offspring from control, unstressed mothers. We asked whether offspring of heat-shocked mothers expressed higher basal levels of the molecular chaperone proteins HSP70 (HSP-1 in *C.* elegans) and HSP90 (DAF-21 in *C.* elegans) which inhibit HSF-1 activation but can also increase thermotolerance, as this might explain the apparently contradictory results of increased stress relilience and decreased stress-induced HSF-1-activity. However, this was also not the case: Western blot analysis using antibodies to endogenous HA tagged HSP-1 (Fig. S4 E) and DAF-21 (Fig. S4 F) showed no differences in protein levels between offspring from stressed mothers and offspring from control, non-stressed mothers. In addiiton, the inducible HSP-70 (C12C8.1) protein levels were also not higher in animals that had experienced early life stress exposure compared to controls (Fig. S4 G), indicating that *hsp-70* mRNA which could have been packaged into soon-to-be-fertilized oocytes was not expressed at measurable levels in animals when they developed into adults and also did not account for their enhanced stress tolerance. Thus, the increased later-life stress resilience of animals that had experienced early-life stress could not be attributed to mechanisms that maintained HSF-1 regulatory regions in a permissive state, nor did it appear to be due to the persistence of gene products that resulted from prior HSF-1 activation. Instead it appeared that a ‘memory’ of early life heat-shock did persist in adult offspring, but resulted in a surprising decrease in the HSF-1 transriptional response to a subsequent heat-shock.

We therefore tested whether MET-2 activity that occurs during normal germline reprogramming was increasing H3K9me2 levels at the transcription start sites or promoter proximal regions of genes induced by HSF-1, due to its germline activity upon heat shock. The persistence of such marks could account for the muted expression of *hsp* genes in these progeny later in life (*41, 42*). RNAi-mediated knockdown of *met-2* in the maternal germline restored the inducible levels of *hsp70* (C12C8.1) in adults that developed from heat-shocked mothers (Fig. S5), suggesting that this may be the case. Therefore, we used ChIP-PCR to more directly measure the levels of H3K9me2 at the 5’-UTR regions of HSF-1 target genes following HSF-1 germline activity (Fig 1C, D; also see Fig. S2 D). For this, we leveraged the 5-minute heat-shock exposure which induces the binding of HSF-1 to *hsp* promoters only in the germline (Fig. S2 A, B). We found that within 2 hours following maternal heat-shock during which time oocytes progress through the maternal germline, H3K9me2 abundance at the promoter proximal 5’-UTR regions of *hsp* genes increased (Fig. 1C). This increase in H3K9me2 at *hsp* genes required germline MET-2, as RNAi-induced knockdown of *met-2*, also only in the maternal germline, abrogated the increase in H3K9me2 at the *hsp* genes (Fig. 1D). The increase in H3K9me2 levels at HSF-1 target genes was also dependent on stress-induced activity of HSF-1 and was not observed in *hsf-1(sy441) I* animals (Fig. 1C). These studies suggested that HSF-1-dependent transcriptional activity that occurred upon heat shock in the maternal germline prompted a MET-2-dependent increase in H3K9me2 at HSF-1 target genes, perhaps in accordance with the role of MET-2 in the epigenetic reprogramming of germline-expressed genes.

We then asked whether the increased H3K9 methylation at heat shock-responsive genes following maternal heat shock reflected an interaction between HSF-1 and MET-2. We used strains where wild type and *hsf-1(sy441) I* mutants were tagged at their endogenous locus with a 3XFLAG tag and *met-2* was tagged at its endogenous locus using 3X HA. We found that HSF-1 directly interacts with MET-2 upon heat shock, as determined by the co-immunoprecipitation of full-length, wild-type HSF-1 with MET-2 (Fig. 1E; compare to the lack of immunoprecipitation with non-specific mouse IgG). MET-2 did not associate with wild-type HSF-1 under control, non-stressed conditions, or with the truncated form of HSF-1 in the *hsf-1 (sy441) I* mutants that lacked the HSF-1 transactivation domain (Fig. 1E). In *C. elegans* and other cells, MET-2 is concentrated at the nuclear periphery throughout development, forming perinuclear heterochromatic foci that associate with genomic regions undergoing transcriptional repression (*43*). We therefore examined whether HSF-1 co-localized with MET-2 at similar structures upon heat-shock. In the absence of heat-shock, both wild type and mutant HSF-1 localize throughout the nucleus in a diffuse pattern. Upon heat-shock or exposure to other stressors, *C. elegans* HSF-1 has been shown to form distinct concentrations within the nucleus, which have been called nuclear stress bodies (nSB), and, albeit of unknown function, are hallmarks of stress and are also formed in some *C. elegans* cells during specific developmental states (*44–47*). Upon heat shock, HSF-1 and MET-2 colocalized at these nSBs, or nuclear foci (Fig 1F, Fig. S6). These foci were distinctly visible in diakinesis oocytes (Fig. 1F), as well as in germ cells in earlier stages of meiosis (early pachytene; Fig. S6 A-D) and in somatic intestinal cells (Fig. S6 E, F) upon a longer (30 minutes) heat shock, but were apparent only in germline cells upon a 5-minute heat shock (Fig. S6 A). In *hsf-1 (sy441) I* mutant animals, HSF-1 did not form nSBs in diakinesis oocytes upon heat-shock (Fig. 1F), and those nSBs formed in pachytene nuclei appeared more disorganized than in wild-type animals (Fig S6 B, C, D). However, MET-2 in *hsf-1(sy441)I* germlines failed to form foci upon heat-shock and remained largely uniform throughout the nuclear periphery and did not co-localize with the mutant HSF-1 nSBs (Fig. 1F, Fig S6 B, C, D).

Consistent with the stress-induced interaction between HSF-1 and MET-2 in diakinesis oocytes resident in the maternal germline, H3K9me2 levels were visibly increased in the oocytes of heat-shocked mothers as assessed by immunostaining with anti-H3K9me2 antibodies (Fig. 2 A, B). Thus, in contrast to the oocytes of unstressed *C. elegans* mothers, where H3K9me2 is usually undetectable, or barely detectable (Fig. 2 A B)(*40*), a significant percent of oocytes in the germline of heat-shocked mothers were enriched for H3K9me2, even when the mothers were only exposed to heat stress for 5 minutes (Fig. 2 A, B). The elevated levels of H3K9me2 persisted for at least up to two hours after the 5-minute heat-shock (Fig. 2 B). Taken together with the increased H3K9me2 levels at HSF-1 target gene 5’-UTR regions, these studies suggested that HSF-1-dependent recruitment of MET-2 in the germline (and other cells) was causing a more global increase in H3K9me2 abundance as was visible in oocytes in the germline of heat-shocked mothers.

**Figure 2:**
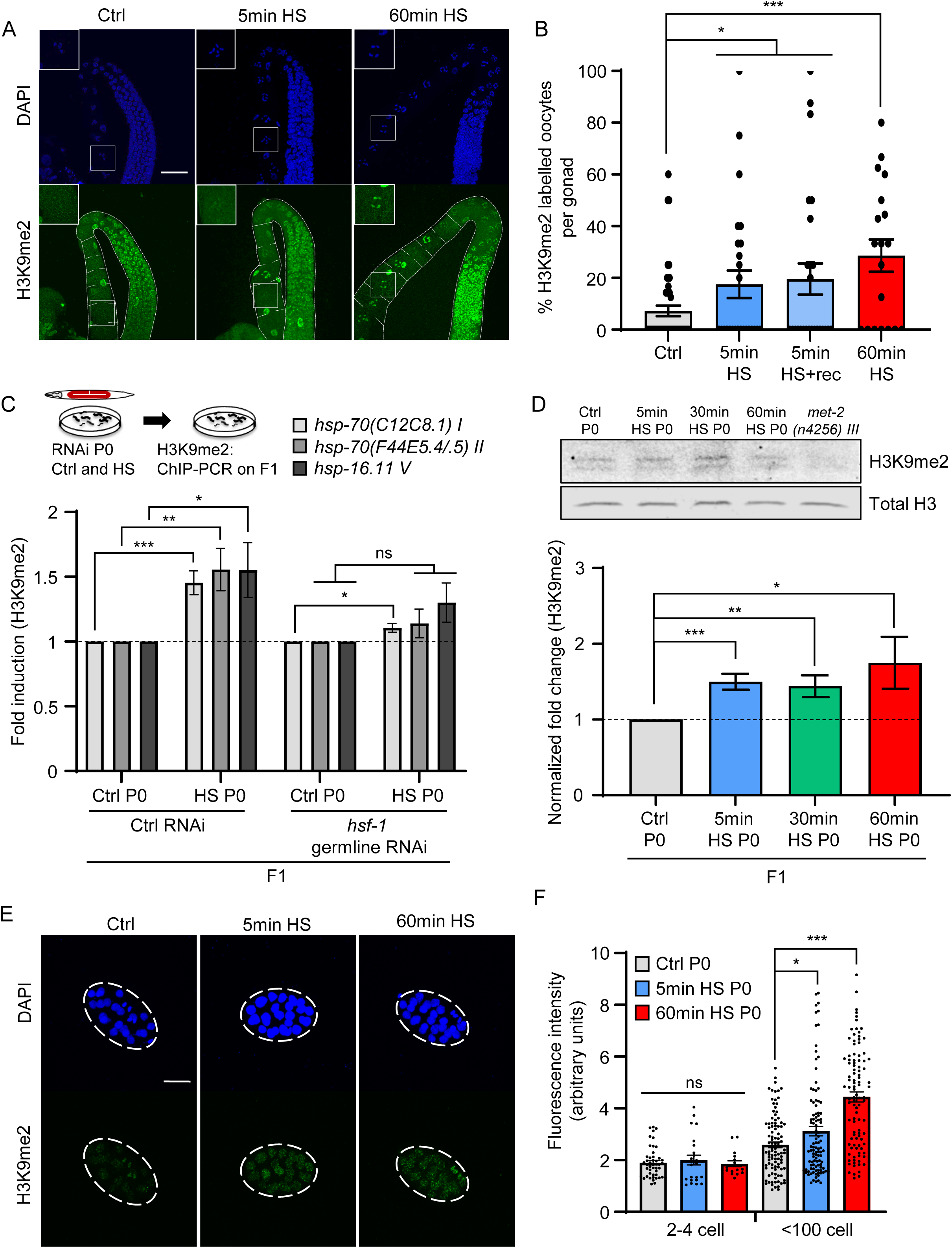
Stress-induced increase in H3K9me2 in oocytes is inherited by progeny. **A,** Representative micrographs showing H3K9me2 staining in projections of confocal z-sections taken through the germline of P0 wild-type animals under control conditions (Ctrl) and upon heat-shock. Heat shock conditions: short 5 minute (5min HS) and long, 60 minute (60min HS) heat shock. Insets: magnified images of diakinesis oocytes. Scale bar:30µm. **B,** Percent H3K9me2 positive diakinesis oocytes per gonad in P0 wild-type animals under control conditions (Ctrl) and upon heat-shock: short 5 minute HS (5minHS), and long 60 minute (60minHS) heat shock. See Materials and Methods for details on quantification. Percent of H3K9me2 positive oocytes present in the P0 maternal germline 2 hours after the mothers were subjected to a short 5 minute heat-shock (5minHS+rec) are also shown. These conditions correspond to conditions where H3K9me2 levels at HSF-1 target genes were assayed (Fig. 1C, D). (n=2-6 experiments; oocytes in 3-15 gonads were quantified per experiment). **C,** H3K9me2 occupancy (relative to control wild-type animals) at the promoter proximal 5’-UTR regions of *hsp-70 (C12C8.1)I, hsp-70 (F44E5.4/.5)II* and *hsp-16.11 V* in day one adult F1 progeny that develop from oocytes of Control P0 mothers, and P0 mothers subjected to a short (5 minute) heat shock. RNAi was used to knockdown *hsf-1* only in the P0 maternal germline. L4440 empty vector was used as the control. (n=5 experiments). **D, Top**: Representative Western blot using anti-H3K9me2 antibody (above) and anti-Histone H3 (below) on F1 progeny that develop from oocytes of P0 mothers subjected to a short (5 minute) or long (30 minute or 60 minute) heat shock. F1 progeny were harvested for western blot analysis after they became day one adults. **Bottom**: H3K9me2 levels quantified relative to total H3 levels and normalized to control animals. *met-2(n4256)III* animals were used to show specificity of antibody. (n=4 experiments). **E,** Representative micrographs showing H3K9me2 staining in F1 embryos from P0 control non-heat shocked mothers and F1 embryos from P0 mothers subjected to a short (5 minute) or long (60 minute) heat shock. Scale bar:15µm. **F,** Fluorescent intensity (arbitrary units) measured from nuclei of 2-4 cell stage F1 embryos and <100 cell stage F1 embryos of control P0 mothers and P0 mothers subjected to a short (5 minute) or long (60 minute) heat shock. (n=2-4 experiments; 17-47 embryonic nuclei for 2-4 cell stage embryo /condition, and 103-105 embryonic nuclei/condition for <100 cell stage embryo were quantified). **B, C, D, F:** Data show Mean ± Standard Error of the Mean *, *p*<0.05; **, *p*< 0.01 ***, *p*<0.001; ns, non-significant; (C;; B, D, F: ANOVA with Tukey’s correction).

For the increased H3K9me2 marks to serve as an epigenetic memory of early life stress exposure, they would have to be inherited by the offspring of heat-shocked parents. This indeed was the case. The increased abundance of H3K9me2 at the 5’-UTR of HSF-1 target *hsp* genes assayed by ChIP-PCR in the mother persisted in adult progeny that developed from the oocytes of these heat-shocked mothers (Fig. 2 C). This increase was not visible in progeny from mothers subjected to RNAi induced knockdown of *hsf-1* in their germline prior to heat-shock (Fig. 2 C). Moreover, consistent with the systemic interaction between HSF-1-and MET-2, and the visible increase in H3K9me2 abundance in oocytes, elevated H3K9me2 levels were apparent even by Western blot analysis in adult progeny that developed from oocytes that had undergone heat-shock while in the maternal germline (Fig. 2 D). To determine whether the increased H3K9me2 levels persisted unaltered though embryogenesis we conducted immunostaining for H3K9me2 on 2-4 cell stage embryos, and embryos in early stages of development dissected from heat-shocked and control mothers soon after fertilization (Fig. 2 E, F). As previously reported (*48, 49*), H3K9me2 is barely detectable at fertilization in embryos under control conditions and increases through embryogenesis to be visible by gastrulation in all embryonic nuclei (*48, 49*). H3K9me2 was still not visible in the 2-4 cell stage embryos generated from heat-shocked oocytes dissected from the maternal germline, but by gastrulation and after, the nuclei of embryos from heat-shocked oocytes had significantly higher levels of H3K9me2 (Fig. 2E, F). Taken together, these data indicate that activated HSF-1 in the maternal germline recruited MET-2 to increase H3K9me2 at select genomic loci in the maternal germline following heat shock, and this increase in H3K9me2 levels was inherited by offspring through some mechanism initiated during the initial heat-shock. Thus, perhaps contrarily, MET-2, which presumably acts to erase germline transcriptional memory was, in the process, reinforcing the memory of transcriptional activation by HSF-1. However, was this activity of MET-2 required for the stress resilience of progeny?

To answer this question, we used RNAi to knock down *met-2* in the maternal germline and assessed the stress resilience of offspring (Fig. 3 A). Knocking down maternal germline *met-2* abolished the intergenerational stress resilience of offspring (Fig. 3 A). As controls, we also examined the effects of knocking down other chromatin modifiers that repress transcription (Fig. 3 A). Knocking down the H3 lysine 9 (H3K9) methyltransferase responsible for H3K9me3 modification of chromatin, SET-25 (*41, 50*), decreased the average stress resilience of progeny but did not have a significant or consistent effect. Knockdown of MES-2 (*51*), the *C. elegans* PRC2 homolog decreased the average stress resilience of offspring after maternal stress exposure, and the knock-down of another SET domain protein SET-1 (*52*) did not alter the protective effects of early life stress heat-shock on offspring stress resilience. We confirmed that MET-2 was required for the thermotolerance of progeny of heat-shocked mothers by also using the deletion mutant *met-2 (n4256) III* (Fig. S7). The rescue of polyQ aggregation in progeny of stressed mothers was also dependent on maternal *met-2*, as RNAi-mediated knockdown of *met-2* in the mother prevented the suppression of protein aggregation in offspring (Fig. 3B). Therefore, it appeared that the maternal activity of MET-2 was indeed required for the stress resilience of progeny and decreasing *met-2* in the maternal germline was sufficient to abolish early life programming of stress resilience.

**Figure 3:**
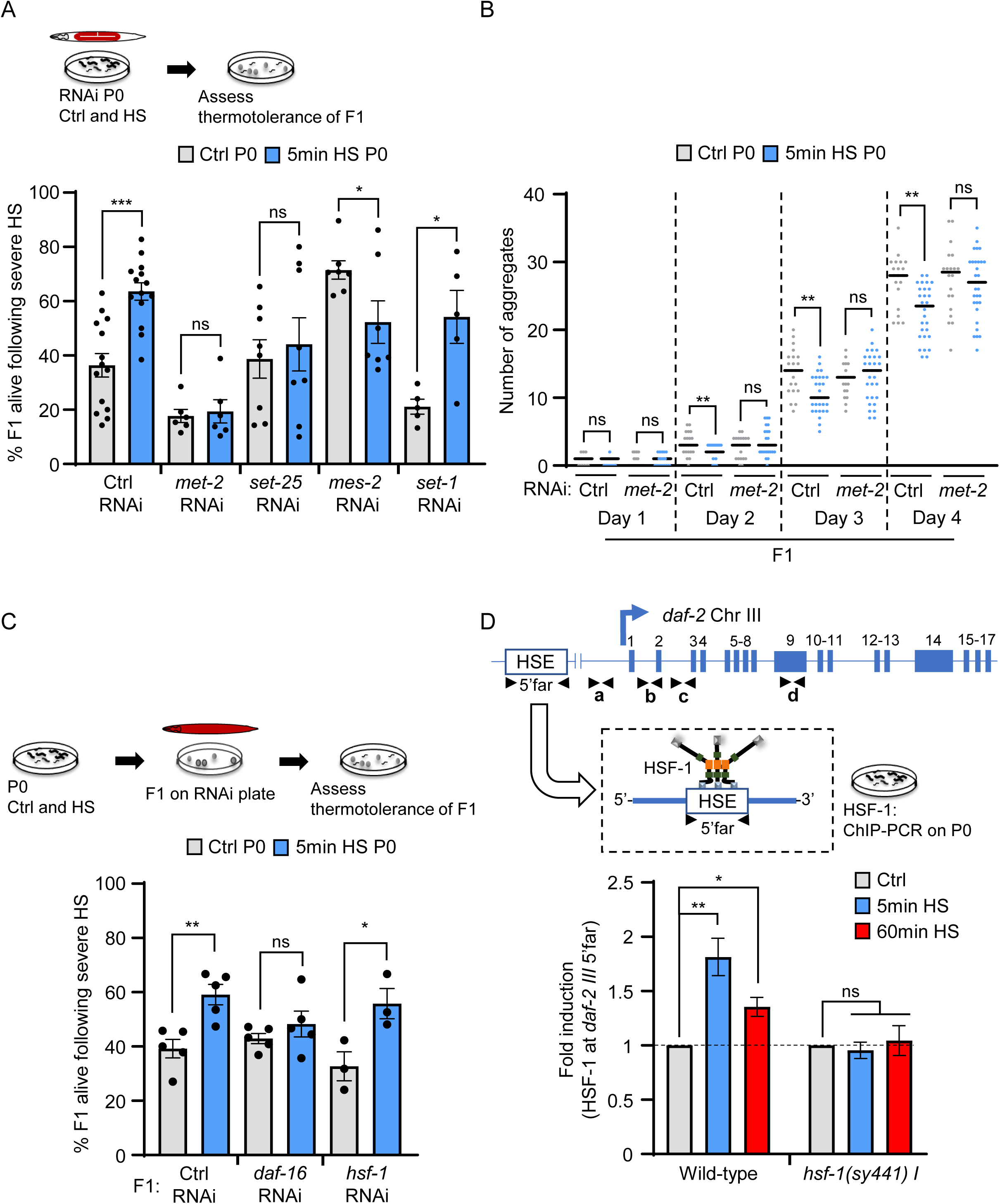
MET-2 activity in the maternal germline is required for progeny stress resilience. **A,** Stress resilience of F1 progeny from control P0 mothers (Ctrl) and P0 mothers subjected to a short (5 minute) heat shock, following the RNAi mediated downregulation of chromatin modifying enzymes (indicated on the X-axis) only in the P0 maternal germline. L4440 empty vector was used as the control. n=4-7 experiments. Each experiment represents thermotolerance of all the progeny laid within the 2-4 hours post-maternal heat shock (average F1 progeny range from 24.1±1 to 28.6±1.4 /experiment). **B,** Number of polyglutamine aggregates scored between day one and day four of adulthood in F1 progeny of control P0 mothers, and P0 mothers subjected to a short (5 minute) heat shock, following the RNAi mediated downreguation of P0 maternal *met-2*. L4440 empty vector was used as the control. (n=3 experiments with 15 animals/ experiment). **C,** Stress resilience of progeny from control P0 mothers, and mothers subjected to a short (5 minute) heat shock, after the F1 progeny were subjected to RNAi mediated knock-down of *daf-16* or *hsf-1*. Note difference from previous experiments: here RNAi was used to downregulate expression in F1 progeny and not in the P0 maternal germline. L4440 empty vector was used as the control. n=3-5 experiments. Each experiment represents thermotolerance of all the progeny laid within 2-4 hours post-heat shock (average F1 progeny scored: 19.1±4.3 to 29±3.3/experiment/condition). **D Top:** Schematic of *daf-2* gene regions analyzed for HSF-1 binding (5’Far; −9071 to −8954). **Bottom**: HSF-1 occupancy at the 5’-Far region (−9071 to −8954) in wild-type animals and *hsf-1(sy441)I* mutants after a short (5 minutes), or long (60 minute) heat shock. (n=3 experiments). **A-D**: Data show Mean ± Standard Error of the Mean *, *p*<0.05; **, *p*< 0.01 ***, *p*<0.001; ns, non-significant; (A, C, D: Unpaired Student’s t-tests and B, paired Student’s t-tests).

These data presented a conundrum: how did HSF-1 and MET-2 activity in the maternal germline provide the progeny with increased stress resilience, while at the same time dampening the stress-inducibility of the very *hsp* genes that are normally needed for surviving stress? One intriguing explanation that suggested itself was that HSF-1-dependent MET-2 recruitment was resulting in the heritable increase in repressive H3K9me2 at some other HSF-1 target(s), whose repression activated stress protection pathways. In *C. elegans*, stress resilience can be largely attributed to the activities of either HSF-1 or the FOXO transcription factor DAF-16 (*36, 53*). Knocking down *hsf-1* only in the progeny of heat-shocked mothers had no effect, consistent with the attenuated induction of *hsps* in these offspring (Fig. 3C, Fig. S8). Knocking down *daf-16* only in progeny from heat-shocked mothers, on the other hand, suppressed their enhanced stress resilience (Fig. 3C, Fig. S8), suggesting that DAF-16 may be activated in the progeny of heat-shocked mothers. Since the protective activity of DAF-16 can be triggered by attenuated signaling of the sole *C. elegans* insulin/IGF-1 receptor, DAF-2, we tested (i) whether HSF-1 indeed bound the 5’UTR region of *daf-2* upon heat shock, (ii) whether HSF-1 and MET-2 activity increased the abundance of H3K9me2 at the *daf-2* locus in heat-shocked mothers which persisted in progeny and (iii) whether DAF-16 was activated in progeny to account for their stress resilience.

Publicly available ChIP-seq data on HSF-1 binding sites in adult *C. elegans* showed an increase in HSF-1 occupancy upon heat-shock at the 5’-UTR of the *daf-2* gene, which contains a putative <u>h</u>eat <u>s</u>hock <u>e</u>lement (HSE)(*35*). We therefore tested whether HSF-1 did indeed bind to this 5’-UTR region of the *daf-2* gene. As determined by ChIP-PCR, HSF-1 occupancy at the *daf-2* 5’-UTR HSE containing region (5’-far; Fig 3D) increased within 5 minutes of heat shock just as it does at its known target *hsp* genes. Also, as seen with the other HSF-1 target genes, this increase did not occur in *hsf-1 (sy441) I* mutants (Fig. 3D) suggesting that HSF-1 binding at the *daf-2* 5’-UTR occurred in response to stress. Interestingly, *daf-2* mRNA levels do not change with heat-shock in wild-type animals or *hsf-1 (sy441) I* mutant animals even after 60 minutes of heat shock (Fig. S9 A). As with other HSF-1 germline targets, H3K9me2 levels also increased across the *daf-2* gene upon heat-shock (Fig. 4A, B). Since multiple isoforms of *daf-2* have been described, we assayed H3K9me2 abundance not only at the 5’-UTR region containing the *daf-2* promoter, but also across the *daf-2* gene body that has been previously shown to be amenable to epigenetic modification (*54*)(Fig. 4A). H3K9me2 levels increased at all these regions in mothers that had been subjected to a 5-minute heat shock and was more elevated in mothers subjected to the longer (60 minutes) heat-shock (Fig. 4A, B). Like the increased H3K9me2 marks at other HSF-1 targets, the elevated levels of H3K9me2 at the *daf-2* gene were inherited by progeny of stressed mothers (Fig. 4C), both when the mothers were subjected to a short (5-minute) and long (30-minute) heat exposure. Also, as with the other HSF-1 targets, this increase of H3K9me2 at the *daf-2* locus in progeny was dependent on HSF-1 activity in the maternal germline: the elevated levels of H3K9me2 at the *daf-2* gene were suppressed when *hsf-1* was knocked down by RNAi in the maternal germline (Fig. 4D). Similarly, the increased H3K9me2 in progeny was also dependent on *met-2* in the maternal germline and was suppressed upon the knockdown of *met-*2 in the maternal germline (Fig. 4D).

**Figure 4:**
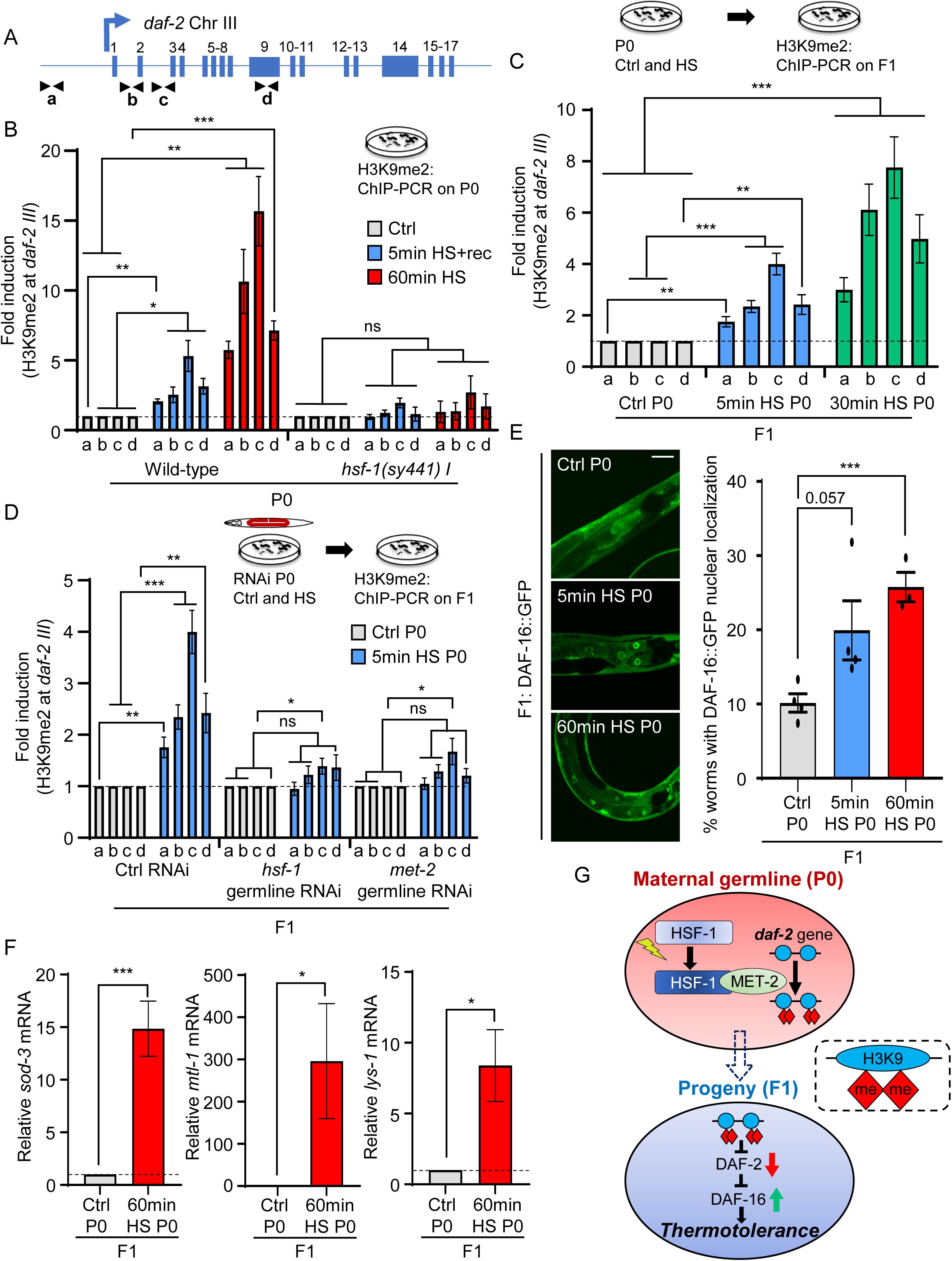
HSF-1 and MET-2 in the maternal germline heritably silence the *daf-2* gene and activate DAF-16 in progeny. **A, Top:** Schematic of *daf-2* gene regions assayed for H3K9me2 occupancy: **a** (−172 to −70), **b** (Intron 1: +3179 to +3279), **c** (Intron 2: +8046 to +8177) and **d** (Exon 9: +15719 to +15834). **B,** H3K9me2 occupancy at regions a, b, c, and d, depicted in **A**, in wild-type and *hsf-1(sy441)I* mutant mothers 2 hours after a 5 minutes heat-shock (5minHS+rec), and immediately after a 60 minute heat shock (n=4 experiments). The longer duration of heat shock provided sufficient time for H3K9me2 accumulation to be detectable in our ChIP-PCR assay, without the need for recovery. **C,** H3K9me2 occupancy at regions a, b, c and d, depicted in **A**, in day one adult F1 progeny from control P0 mothers and P0 mothers subjected to a short (5 minute), or long (30 minute) heat shock (n=4 experiments). **D,** H3K9me2 occupancy at regions a, b, c and d, depicted in **A**, in day one adult F1 progeny from control P0 mothers, and P0 mothers subjected to a short (5 minute), heat shock following RNAi mediated knockdown of *hsf-1* or *met-2* in the P0 maternal germline. (n=5 experiments). **E**, **Left:** Representative micrographs showing DAF-16::GFP localization in day one adult F1 progeny from control P0 mothers (Ctrl), and P0 mothers subjected to a short (5 minute) or long (60 minute) heat shock. Scale bar:30µm. **Right:** Percent F1 progeny that display DAF-16::GFP nuclear localization as day one adults. (n=3-4 experiments, 20-35 progeny scored per experiment). **F,** Average *sod-3, mtl-1 and lys-1* mRNA levels in day one adult F1 progeny from control P0 mothers (Ctrl), and P0 mothers subjected to a 60 minute heat shock. mRNA levels were determined relative to *pmp-3*, and normalized to that in F1 progeny from control P0 mothers (n=4 experiments; 30 animals from 15 mothers/experiment). **G,** Model: germline activity of HSF-1 induces a MET-2-dependent increase in H3K9me2 at the *daf-2* gene; this repression of *daf-2* is inherited by F1 progeny of the next generation and DAF-16 is activated to confer thermotolerance.

Consistent with the increase in H3K9me2 at *daf-2*, *daf-2* mRNA levels were decreased in the progeny of heat-shocked mothers (Fig.S9 B). This decrease in *daf-2* expression was functional and DAF-16 was activated in the F1 progeny that had been subjected to heat-shock as oocytes in the maternal germline, even when these progeny became adults. This was assayed by the increased nuclear localization of a DAF-16::GFP transgenic protein in F1 progeny of stressed mothers grown under normal control conditions (Fig. 4E) and confirmed by the increased expression of known DAF-16 target genes in these F1 progeny (Fig. 4F). Thus, maternal stress exposure of *C. elegans* activated HSF-1 in the germline and induced a MET-2 dependent increase in H3K9me2 marks at the *daf-2* locus, to program the activity of the Insulin-like signaling pathway and stress-resiliency in the next generation (Fig. 4G).

In summary, these studies demonstrate a novel mechanism whereby in response to stress, the germline activity of the conserved transcription factor HSF-1 recruits the repressive histone H3K9 SET domain methyltransferase MET-2 to HSF-1 target genes, including to the single insulin receptor *daf-2*, to increase heterochromatin and repress their subsequent expression. We propose that this occurs at sites in the nucleus similar to previously described heterochromatin foci formed by MET-2 during normal development, but are likely to contain new genomic loci recruited by HSF-1 and MET-2 for silencing. While this could arguably be similar to the ‘normal’ epigenetic programming mechanisms that exist to reset the transcriptional memory associated with germline gene expression, it results in the heritable increase in epigenetic marks that enhance the stress resilience of offspring, likely providing them with a survival advantage if the environment remained suboptimal. We therefore hypothesize that this mechanism may represent a novel and consequential function of HSF-1 in the establishment of an epigenetic memory of stress. In addition to functioning as a transcription factor, HSF-1 binds to additional sites in chromatin in all organisms, the function of which, in most cases, is not apparent (*55, 56*). In *C. elegans* DAF-2 insulin-like signaling pathway regulates lifespan, development, neuronal function, stress resistance, immunity, and multiple aspects of metabolism (*54, 57–69*). Therefore, the binding of HSF-1 to the *daf-2* promoter to repress *daf-2* in response to early life stress exposure is likely to cause long-term changes in gene expression, physiology and functioning of the organism. While much remains to be understood, considering the dysregulation of both HSF1 and the mammalian MET-2 homolog, SETDB1 (SET Domain Bifurcated Histone Lysine Methyltransferase 1) in cancer, and the alterations of H3K9me2 that occur at chromatin in neurodegenerative disease of aging, we speculate that HSF-1-mediated heterochromatinization through MET-2/SETDB1 recruitment, as seen here, could be activated in humans in response to environmental toxins, pathogens or starvation, and result in a similar cascade of epigenetic changes during perinatal development to affect the long-term physiology of offspring.

## Acknowledgements

We thank the members of V.P. laboratory, Dr. Sarit Smolikove, Dr. Josep Comeron, Dr. Jian Li, Dr. Tali Gidalevitz and Dr. Yair Argon for their helpful comments and Dr. Rachel Reichman and Dr. Smolikove for advice with CRISPR/Cas9. Nematode strains were provided by the Caenorhabditis Genetics Center (CGC) (funded by the NIH Infrastructure Programs P40 OD010440). This work was supported by NIH R01 AG 050653 (V.P.).

---

**Figure S1:**
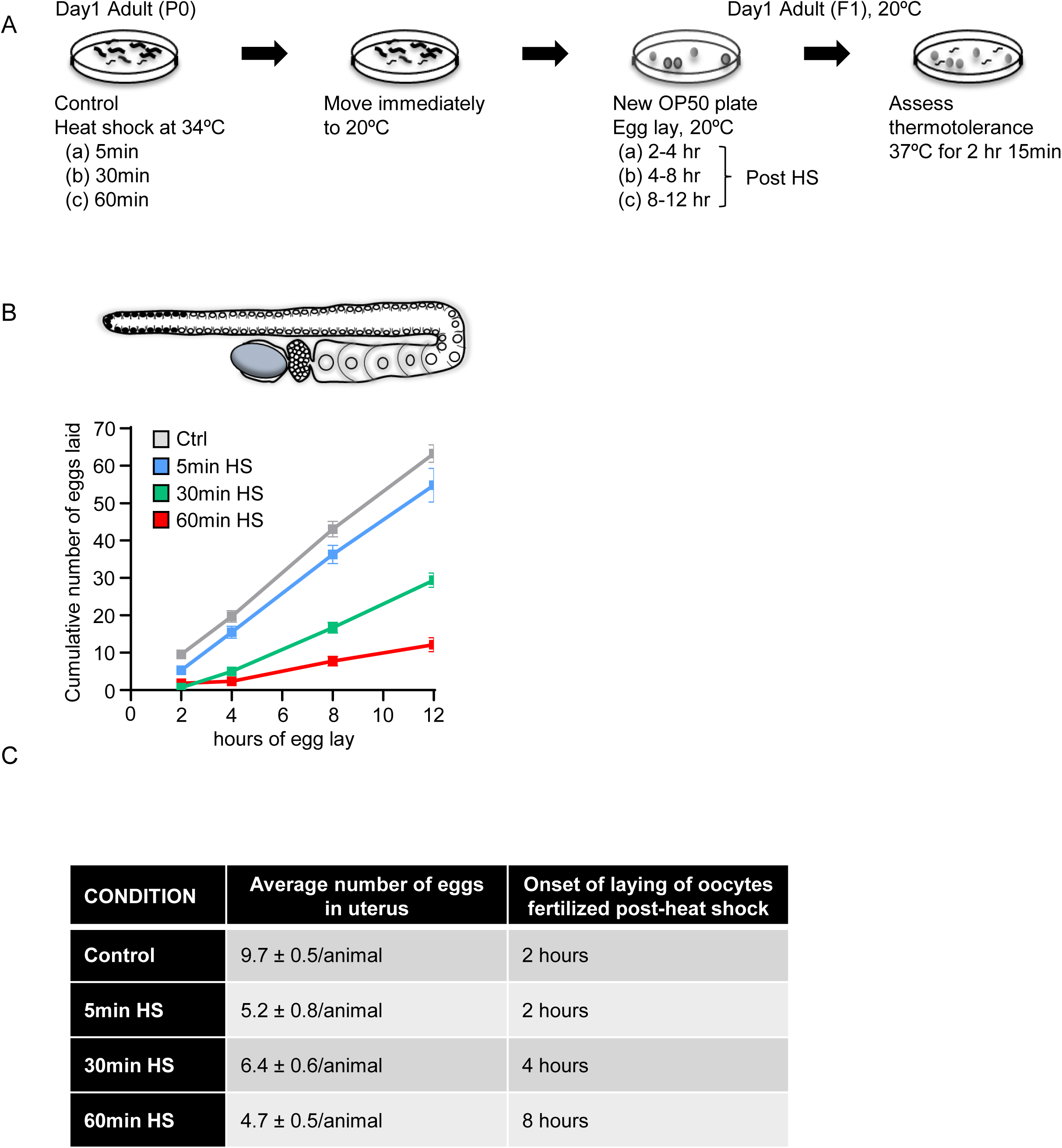
Experimental protocol to activate HSF-1 in germ cells of *C. elegans* and examine the later-life phenotypes of offspring generated from these germ cells. **A**, Schematic of experimental procedure. Adult, day-one-old P0 mothers were exposed to 34°C for short (5 minute) or long (30 minute or 60 minute) durations. Following heat shock, P0 mothers were moved to control 20°C conditions. Embryos laid 2-4 hours, 4-8 hours or 8-12 hours following a 5 minute, 30 minute or 60 minute maternal heat shock respectively (F1) were collected on separate plates. These embryos were estimated to be generated by the fertilization of the diakinesis oocytes resident in the maternal germline during heat-shock, but fertilized and laid at 20°C (see **B** and **C**), and were used for experiments. Control animals were age matched. F1 progeny were allowed to become day-one adults at 20°C, and subjected to a severe heat shock of 37°C for 2 hours, 15 minutes (chosen after prior experiments to titrate survival of control animals to <∼50% and prevent ceiling effects). **B,** Y-axis: Total cumulative number of embryos laid by control P0 mothers and P0 mothers subjected to 5 minutes, 30 minutes and 60 minutes heat shock, at different times (X-axis) following heat shock. These numbers were calculated by counting the number of embryos laid every 2 hours for upto 12 hours. **C**, **Table Column 1**: P0 mothers’ treatment conditions. **Table Column 2**: Average numbers of already fertilized eggs in the uterus of mothers under control conditions, and <u>immediately after</u> heat-shock exposure. The number of unfertilized oocytes in all animals (8.6 ± 0.5/animal). **Table Column 3**: The time post-heat shock at which oocytes resident in the maternal germline during heat shock will start being laid post-fertilization, as calculated from the information in **B**, and **Column 2**.

**Figure S2:**
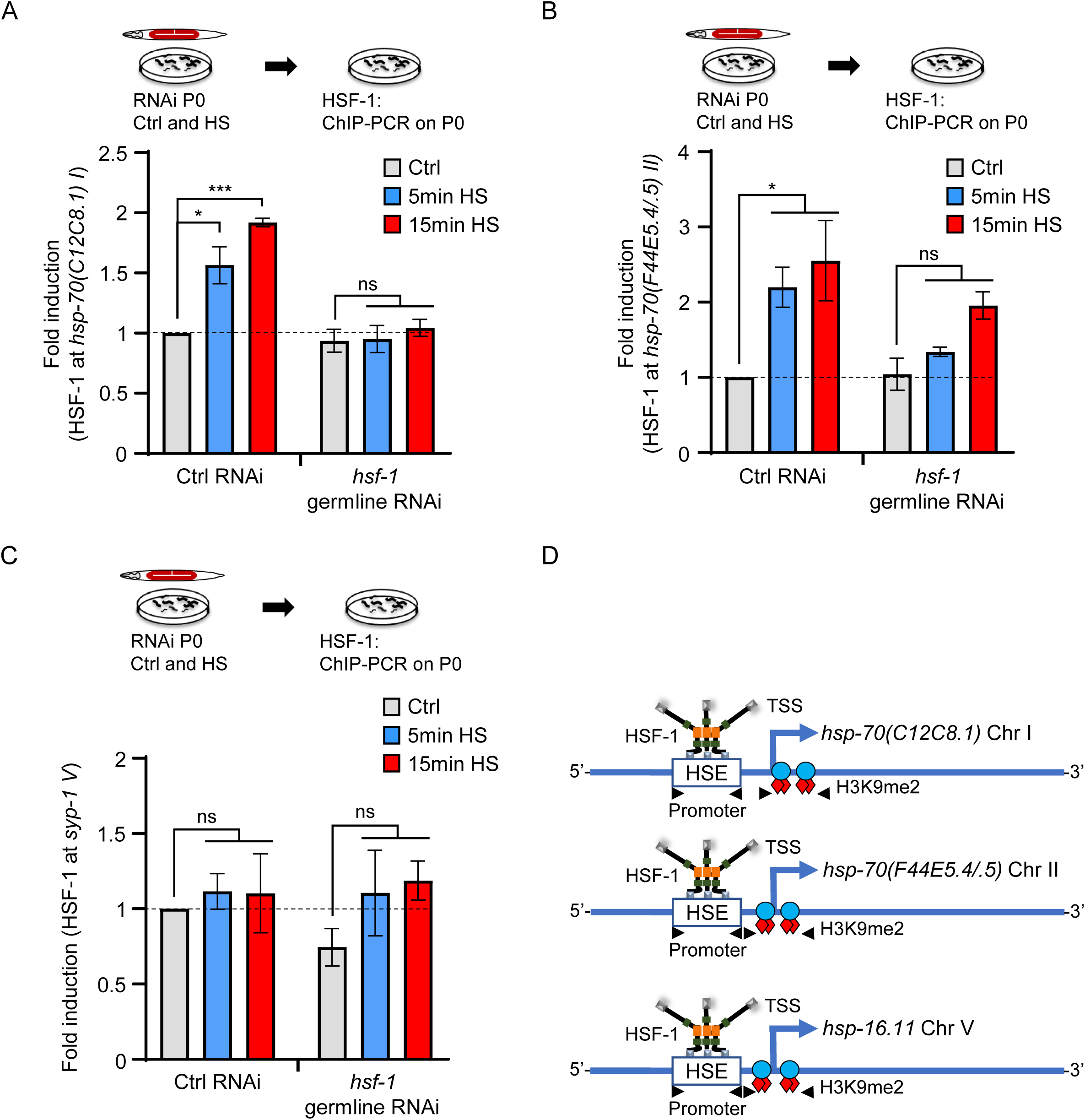
HSF-1 is activated in the germline upon heat shock. **A,** HSF-1 occupancy at the Promoter regions of *hsp-70 (C12C8.1) I* under control conditions and upon a short (5 minute), and long (15 minute) heat-shock was determined in wild-type animals and animals where *hsf-1* was knocked down in the germline (n=6 experiments). Note: HSF-1 binding is abolished upon both short and long heat shock (*p*<0.05 and *p*< 0.01, Unpaired student’s t-test) when HSF-1 is knocked down only in the germline, even though occupancy at *hsp-70* regions is assessed in chromatin purified from whole animals. **B,** HSF-1 occupancy at the Promoter regions of *hsp-70 (F44E5.4/.5) II* under control conditions and upon a short (5 minute), and long (15 minute) heat-shock was determined in wild-type animals and animals where *hsf-1* was knocked down in the germline (n=6 experiments). Note: HSF-1 binding is significantly decreased upon the short heat shock (*p*<0.05, Unpaired student’s t-test) even when HSF-1 is only knocked down in the germline. **C,** Specificity of HSF-1 binding in **A**, and **B**, verified by assessing HSF-1 occupancy at the *syp-1* promoter which does not possess a HSE, and is not known to bind HSF-1. **D,** Schematic of *hsp-70 (C12C8.1) I*, *hsp-70 (F44E5.4/.5) II*, and *hsp-16.11 V* promoter and 5’-UTR regions analyzed for HSF-1 and H3K9me2 binding. For exact positions see Materials and Methods. **A, B, C:** Data show Mean ± Standard Error of the Mean. Data in all experiments are normalized to values from control (non-heat shocked) wild-type animals. *, *p*<0.05; ***, *p*<0.001; (Unpaired Student’s t-test). ns, non-significant.

**Figure S3:**
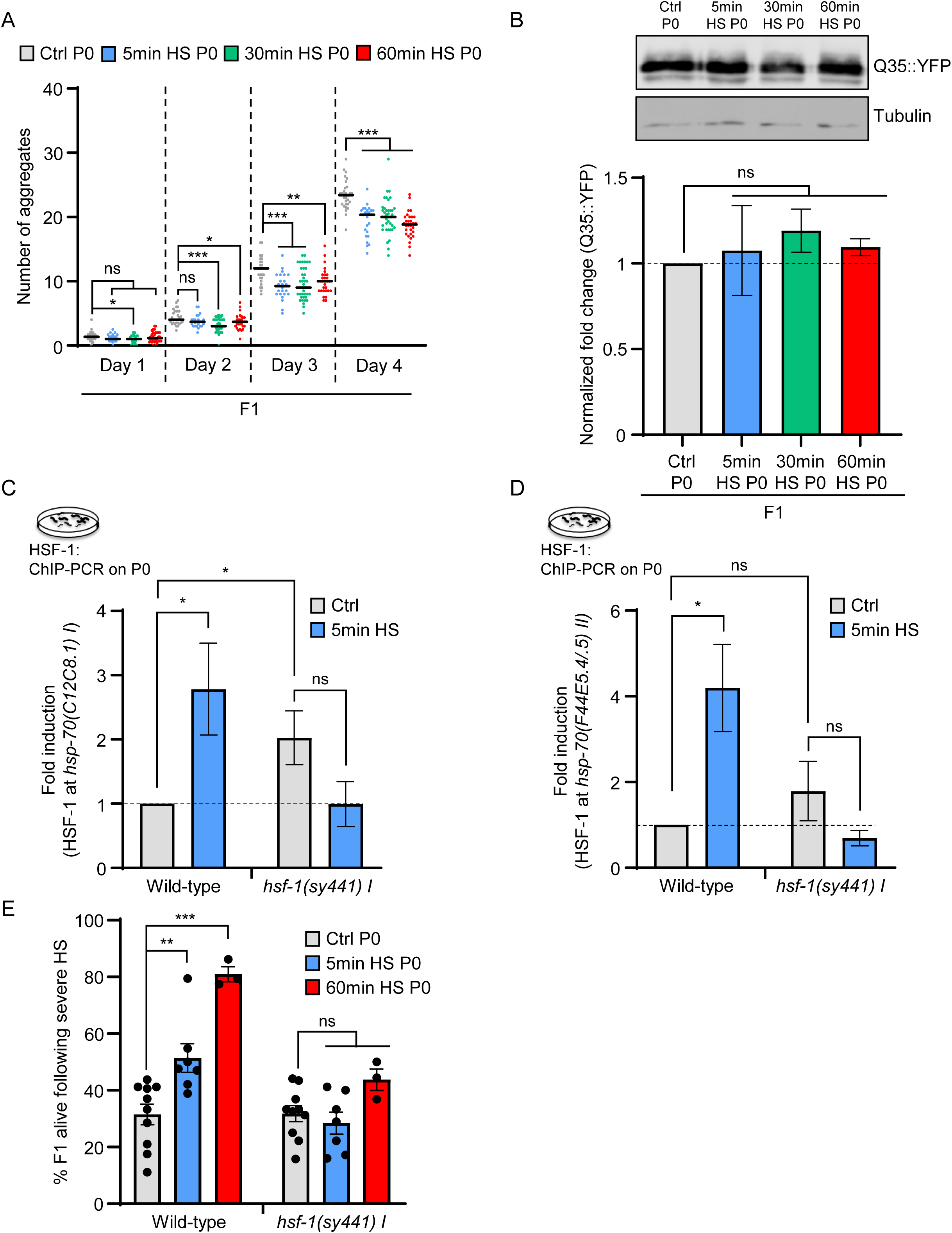
Maternal HSF-1 protects progeny stress resilience and proteostasis. **A,** Number of polyglutamine aggregates scored from day one to day four of adulthood in F1 progeny of control P0 mothers, and P0 mothers subjected to a short (5 minute) heat shock and long (30 minute or 60 minute) heat shock. (n=3 experiments with 25 animals/ experiment). **B, Top:** Representative Western blot showing polyglutamine protein levels in day one adult F1 progeny of control P0 mothers, and P0 mothers subjected to a short (5 minute) heat shock and long (30 minute or 60 minute) heat shock (above). Tubulin was used as the loading control (below). Day one adults were chosen as the majority of polyglutamine is still soluble at that age. **Bottom:** Quantitation of polyglutamine protein levels in day one adult F1 progeny. **C,** HSF-1 occupancy at the Promoter regions of *hsp-70 (C12C8.1)I* under control conditions and upon a short (5 minute) heat-shock in wild-type animals and *hsf-1 (sy441)I* mutants (n=6 experiments). **D,** HSF-1 occupancy at the Promoter regions of *hsp-70 (F44E5.4/.5)II* under control conditions and upon a short (5 minute) heat-shock in wild-type animals and *hsf-1 (sy441)I* mutants (n=6 experiments). **E,** Stress resilience of wild-type and *hsf-1 (sy441)I* mutant F1 progeny of control P0 mothers and P0 mothers subjected to a short (5 minute) or long (60 minute) heat shock. (n=3-9 experiments).Data show Mean ± Standard Error of the Mean. *, *p*<0.05; **, *p*< 0.01 ***, *p*<0.001; (A, ANOVA with Tukey’s correction; B-E, Unpaired Student’s t-test). ns, non-significant.

**Figure S4:**
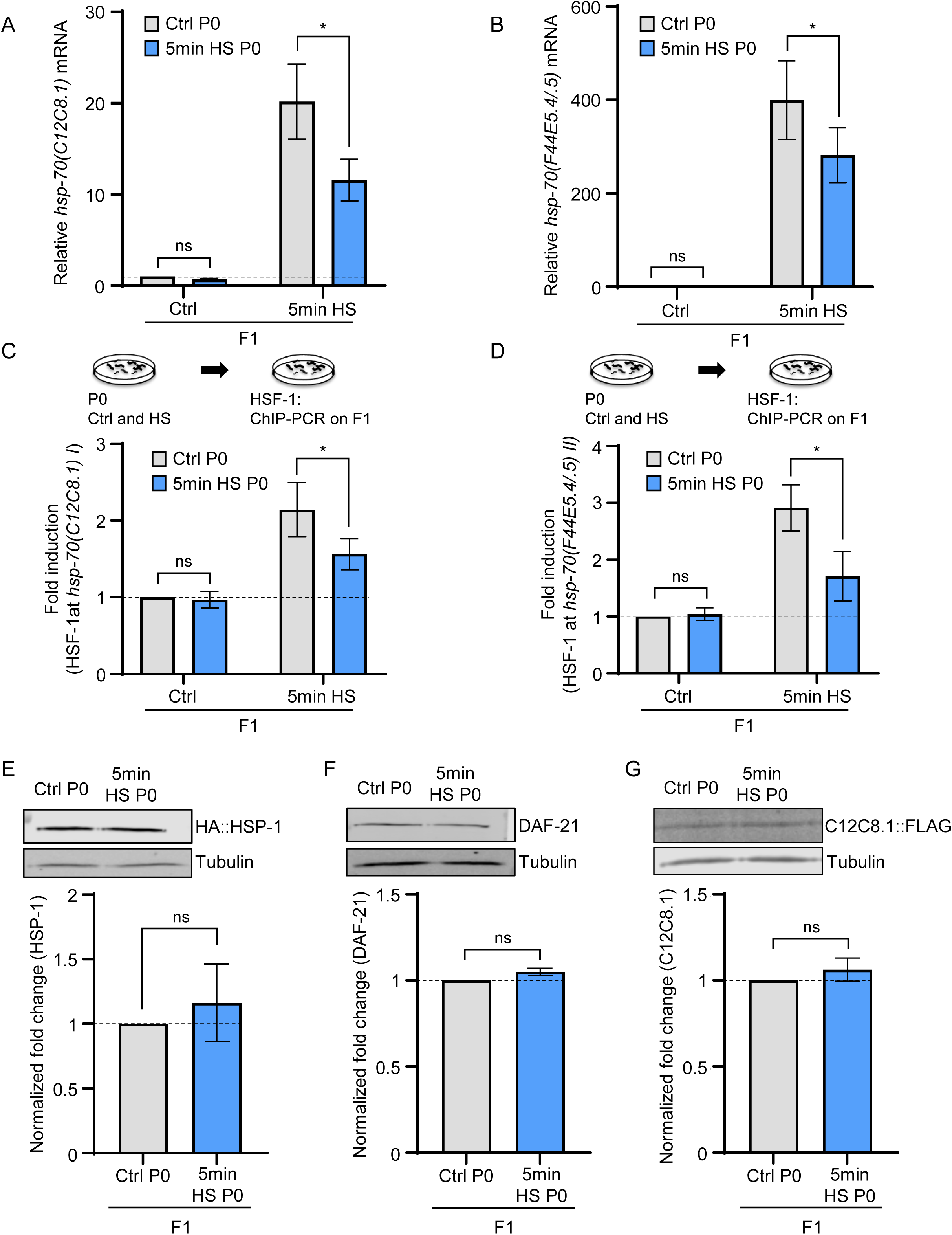
HSF-1 activity in the mother results in a decrease in the inducible heat-shock response in progeny. **A,** Average *hsp70* (*C12C8.1*) mRNA levels and **B,** average *hsp70* (*F44E5.4/.5*) mRNA levels in F1 progeny of control P0 mothers, and F1 progeny of P0 mothers subjected to a 5 minute heat-shock. mRNA levels in F1s were assessed under control conditions and after they were also subjected to heat-shock (5 minutes). mRNA levels were normalized to that in F1 progeny of control non-heat shocked, P0 mothers. (n=4 experiments). **A, B** mRNA levels were determined relative to *pmp-3*, and normalized to that in F1 progeny from control P0 mothers. **C,** HSF-1 occupancy at the promoter region of *hsp-70 (C12C8.1) I,* and, **D,** at the promoter region of *hsp-70 (F44E5.4/.5) II* in F1 progeny of P0 control mothers, and F1 progeny of P0 mothers subjected to a short (5 minute) heat-shock. HSF-1 occupancy in F1s was assessed under control conditions and after they were also subjected to heat-shock (5 minutes). HSF-1 occupancy was normalized to that in F1 progeny of control non-heat shocked, P0 mothers (n=4 experiments). **E, Top:** Representative Western blot showing HSP-1 protein levels (above) in day one adult F1 progeny of control P0 mothers, and P0 mothers subjected to a short (5 minute) heat shock. Tubulin was used as the loading control (below). **Bottom:** Quantitation of HSP-1 protein levels in day one adult F1 progeny. (n=3 experiments). **F, Top:** Representative Western blot showing DAF-21/HSP90 protein levels (above) in day one adult F1 progeny of control P0 mothers, and P0 mothers subjected to a short (5 minute) heat shock. Tubulin was used as the loading control (below). **Bottom:** Quantitation of DAF-21 protein levels in day one adult F1 progeny. (n=3 experiments). **G, Top:** Representative Western blot showing HSP-70 (C12C8.1) protein levels (above) in day one adult F1 progeny of control P0 mothers, and P0 mothers subjected to a short (5 minute) heat shock. Tubulin was used as the loading control (below). **Bottom:** Quantitation of HSP-70 (C12C8.1) protein levels in day one adult F1 progeny. (n=2 experiments). Data show Mean ± Standard Error of the Mean. *, *p*<0.05; (Paired Student’s t-test). ns, non-significant.

**Figure S5:**
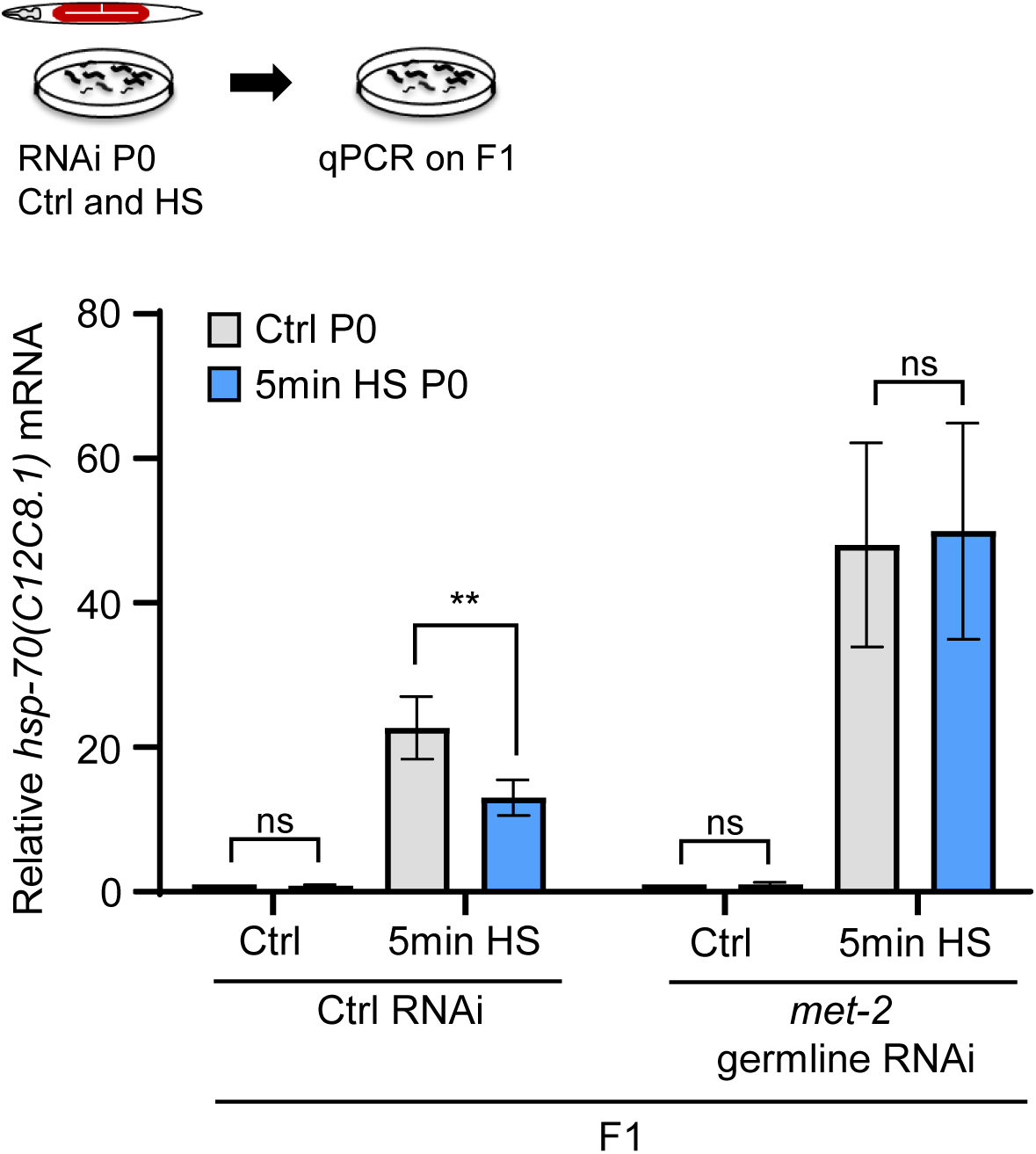
Maternal germline *met-2* is reponsible for the diminished expression of *hsp* genes in F1 progeny of heat shocked mothers. Average *hsp70* (*C12C8.1*) mRNA levels in F1 progeny of control P0 mothers, and F1 progeny of P0 mothers subjected to a 5 minute heat-shock, after control (L4440) RNAi and *met-2* RNAi in the maternal germline. mRNA levels were normalized to that in F1 progeny of control non-heat shocked, P0 mothers. (n=3 experiments). mRNA levels were determined relative to *pmp-3*, and normalized to that in F1 progeny from control P0 mothers. Data show Mean ± Standard Error of the Mean. **, *p*< 0.01; (Paired Student’s t-test). ns, non-significant.

**Figure S6:**
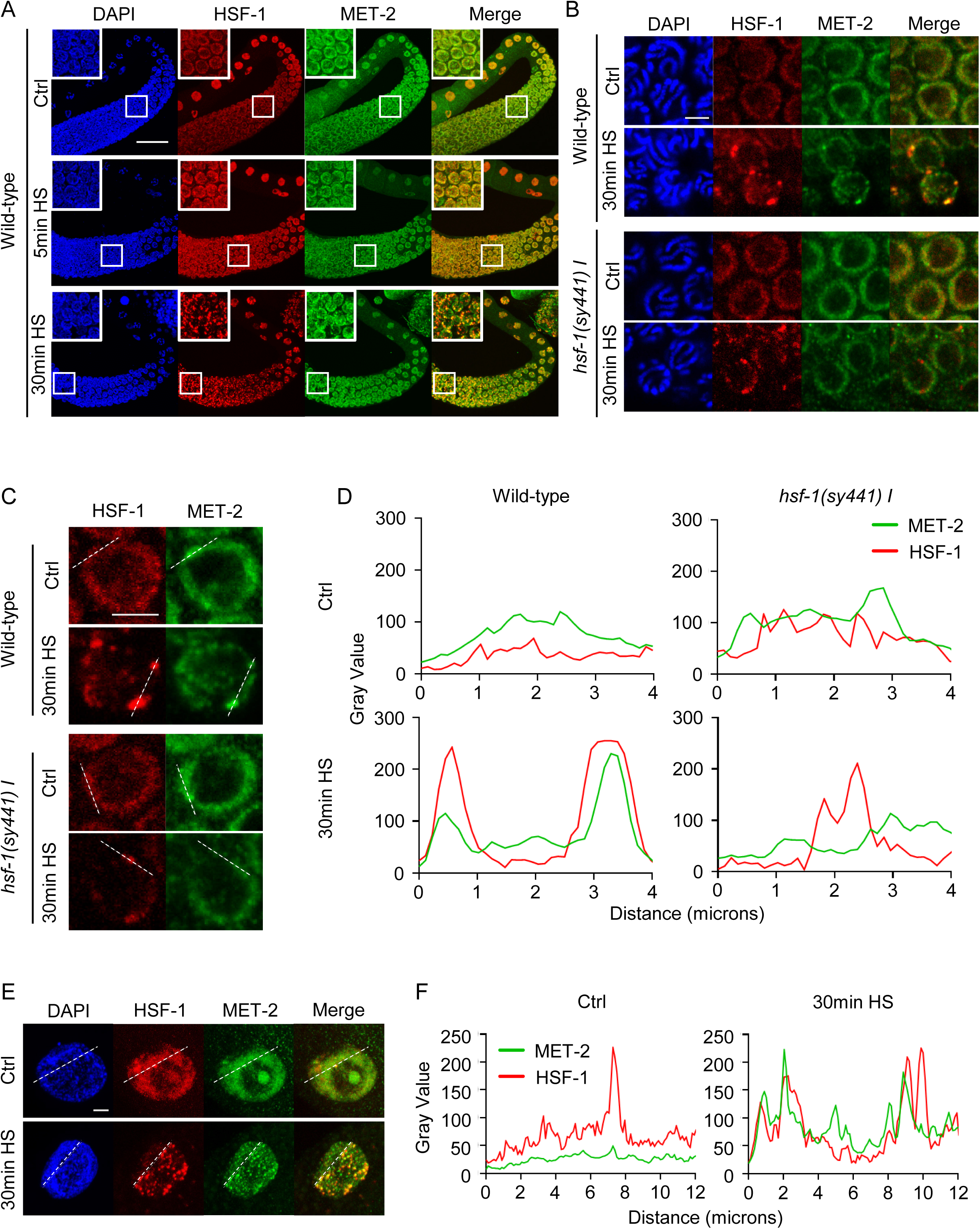
Wild-type HSF-1, but not the truncated HSF-1 in *hsf-1 (sy441) I* mutant animals colocalizes with MET-2 in nuclear stress bodies (nSBs) upon heat shock. **A,** Representative micrographs showing projections of confocal z-sections through dissected germlines of wild-type animals under control conditions and upon short heat-shock (5 minutes) and long heat shock (30 minutes). Insets show maginified regions from pachytene. Scale bar:30µm**. B,** Representative micrographs of a confocal z-section through pachytene germ cells in wild-type animals and *hsf-1(sy441) I*, under control conditions and upon 30 minute heat-shock, Scale bar:3µm. **C,** Magnified image of one nucleus from **B.** Scale bar:3µm. **D,** Intensity profile graphs of HSF-1 colocalization with MET-2 in nuclei from **C**. Line scan graphs were generated by plotting the immunofluorescence intensity along the freely positioned line at the periphery of the nucleus in **C**. **E,** Representative micrographs showing projections of confocal z-sections through intestinal nuclei in wild-type animals under control conditions and upon 30 minute heat-shock. Scale bar:3µm. **F,** Intensity profile graphs of HSF-1 colocalization with MET-2 (plotted as immunofluorescence intensity along the freely positioned line) of nuclei in **E**. **A, B,C, E:** Red: HSF-1. Green: MET-2. DAPI: DNA.

**Figure S7:**
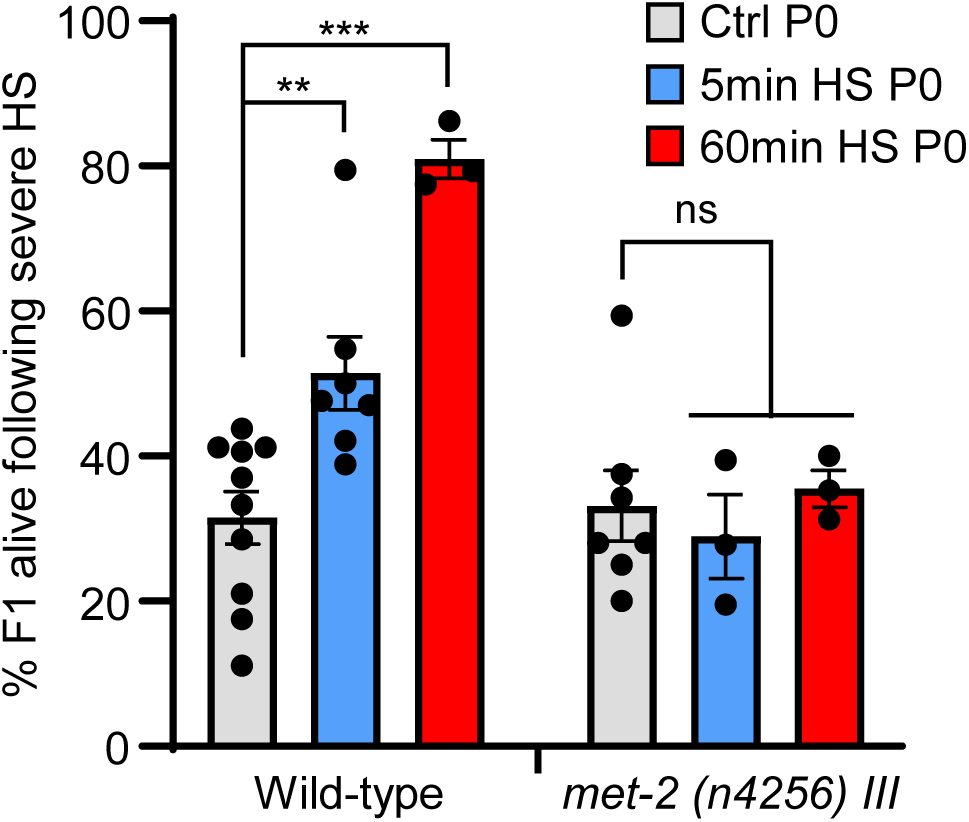
MET-2 is required for the increased stress reslience of F1 progeny of heat shocked mothers. Stress resilience of F1 progeny of wild-type and *met-2 (n4256)III* mutants. F1 progeny are from control P0 mothers and P0 mothers subjected to a short (5 minute) or long (60 minute) heat shock. (n=3-10 experiments). Data show Mean ± Standard Error of the Mean. **, *p*< 0.01 ***, *p*<0.001; (Unpaired Student’s t-test). ns, non-significant.

**Figure S8:**
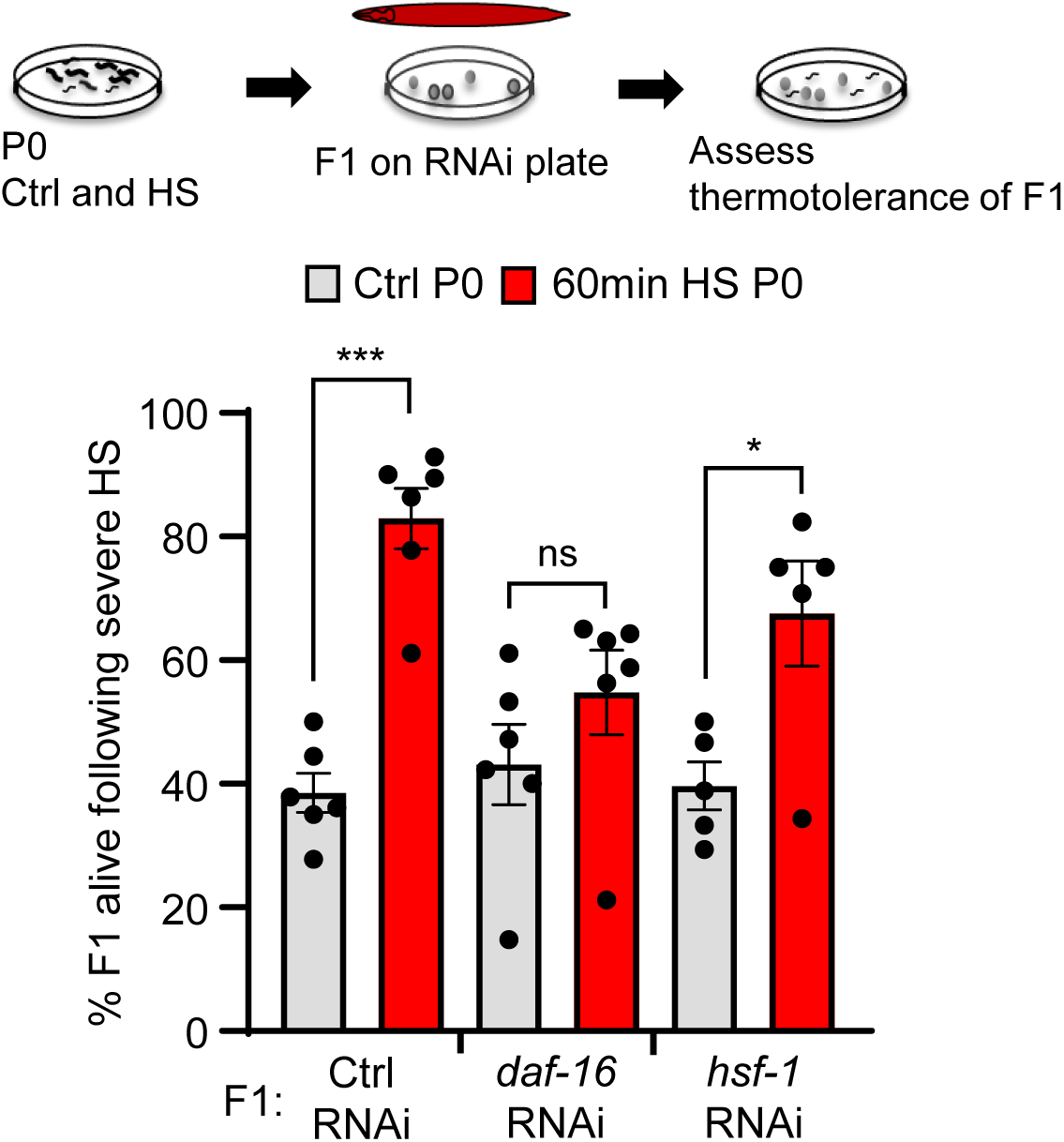
DAF-16 activity, but not HSF-1 activity is required for stress resilience of F1 progeny of heat-shocked mothers. Stress resilience of progeny from control P0 mothers and mothers subjected to a long (60 minute) heat shock, after the F1 progeny were subjected to RNAi mediated knock-down of *daf-16* or *hsf-1*. Note the difference from previous experiments: here RNAi was used to downregulate expression in F1 progeny and not in the P0 maternal germline. L4440 empty vector was used as the control. n=5-6 experiments. Each experiment represents thermotolerance of all the progeny laid within 8-12 hours post-heat shock (average F1 progeny scored: 22±3.8 to 30.1±5.2/experiment/condition). Data show Mean ± Standard Error of the Mean. *, *p*<0.05; ***, *p*<0.001; (Unpaired Student’s t-test). ns, non-significant.

**Figure S9:**
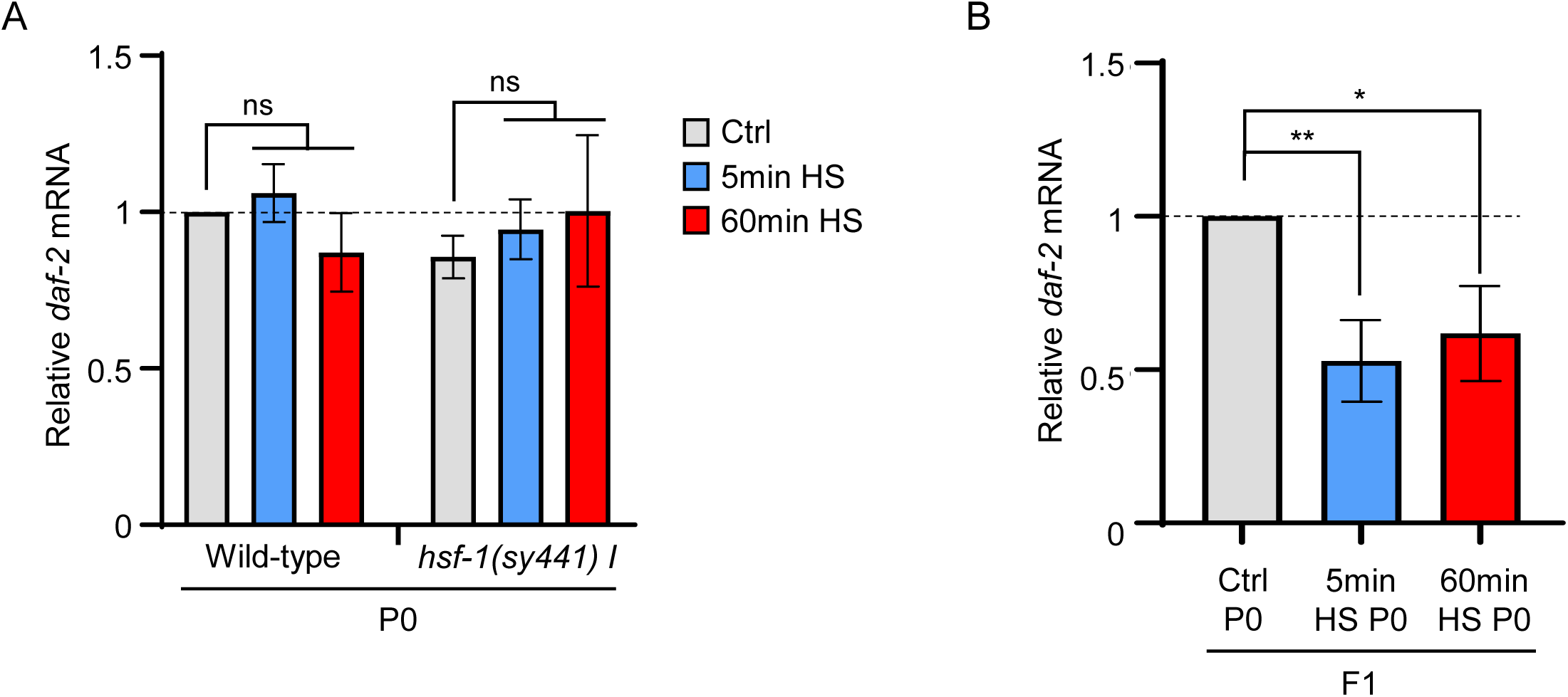
*daf-2* mRNA is not induced by heat shock, but is downregulated in F1 progeny of heat-shocked mothers. **A,** Average *daf-2* mRNA levels in wild-type and *hsf-1(sy441)I* animals under control conditions and upon a short (5 minute) and long (60 minute) heat-shock. mRNA levels were normalized to that in control non-heat shocked, wild-type animals. (n=3 experiments). **B,** Average *daf-2* mRNA levels in F1 progeny of control, non-heat shocked P0 mothers, and F1 progeny of P0 mothers subjected to a short (5 minute) and long (60 minute) heat shock. mRNA levels were normalized to that in that in F1 progeny from control non-heat shocked mothers. **A, B,** mRNA levels were determined relative to *pmp-3*. Data show Mean ± Standard Error of the Mean. *, *p*<0.05; **, *p*< 0.01; (Unpaired Student’s t-test). ns, non-significant.

## *C. elegans* strains

*C. elegans* used in this study are listed below. Strains were procured from Caenorhabditis Genetics Center (CGC, Twin Cities, MN), generated in the Prahlad Lab, or generated by SunyBiotech (Suzhou, Jiangsu, China 215028).

| Strain Name | Genotype | Source |
| --- | --- | --- |
| Wild-type | Bristol N2 | CGC |
| PS3551 | *hsf-1(sy441)I* | CGC |
| AM140 | *rmIs132 [unc-54p::Q35::YFP]* | CGC |
| DCL569 | *mkcSi13 II; rde-1(mkc36) V* | CGC |
| MT13293 | *met-2(n4256) III* | CGC |
| TJ356 | *zIs356 [daf-16p::daf-16a/b::GFP + rol-6(su1006)] IV* | CGC |
| HSF-1::FLAG | *hsf-1::3X flag I* | Prahlad lab (1) |
| FLAG::HSF-1 | *3X flag::hsf-1( I)* | Prahlad lab |
| FLAG::HSF-1(sy441) | *3Xflag:: hsf-1(sy441)I* | Prahlad lab |
| PHX2467 | *hsp-1(syb2467)*<br>(HA::HSP-1) | Prahlad lab / SunyBiotech |
| PHX2141 | *hsp-70(syb2141)*<br>(C12C8.1::FLAG) | Prahlad lab/ SunyBiotech |
| PHX3574 | *met-2(syb3574)*<br>(MET-2::HA) | Prahlad lab/ SunyBiotech |

## Generation of CRISPR strains

CRISPR/Cas9 was used to create following *C. elegans* strains by adding different tags at endogenous loci:

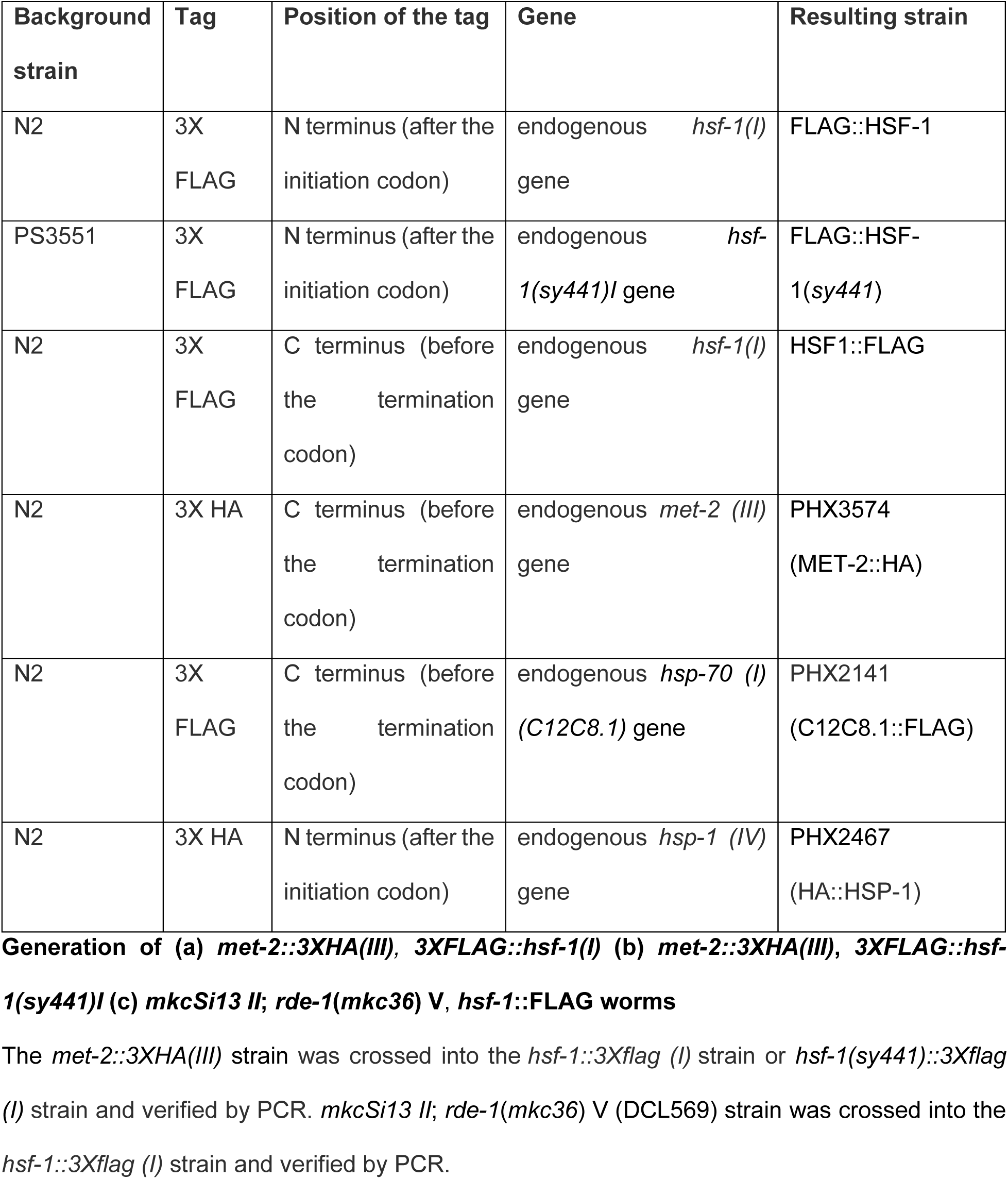

## Growth conditions of *C. elegans* strains

All strains were grown and maintained at 20°C. Animals were grown and maintained at low densities in incubators under standard conditions by passaging 8–10 L4s onto nematode growth media (NGM) plates and, 4 days later, picking L4 animals onto fresh plates for experiments. Animals were fed *Escherichia coli* OP50 obtained from Caenorhabditis Genetics Center (CGC) that were seeded onto culture plates 2 days before use. The NGM plates were standardized by pouring 8.9 ml of liquid NGM per 60 mm plate. Plates had an average weight of 13.5 ± 0.2 g. Any plates that varied from these measurements were discarded. Laboratory temperature was maintained at 20°C to 22°C and carefully monitored throughout the experimental procedures. All animals included in the experiments, unless otherwise stated, were one-day-old hermaphrodites that were age-matched either by (a) bleaching and starting the experiment after 75–78 hrs. or (b) picking as L4 juveniles 24 to 26 hrs. before the start of the experiment.

## Heat shock of worms

The heat shock protocol described in Das et al., 2020 (*1*) was used. Briefly, NGM plates (8.9 ml liquid NGM/plate, weight 13.5 ± 0.2 g) were seeded with 250-300 µl OP50 culture in the center and allowed to dry at room temperature for 48 hr. Either L4 hermaphrodites were passaged onto these plates or worms were bleach-hatched onto these plates and allowed to grow to Day1 adults. All heat shock experiments were performed with one-day-old gravid animals. To induce heat shock response in *C. elegans*, NGM plates containing one-day-old animals were parafilmed and immersed in water bath (product no. ITEMP 4100 H21P 115V, Fischer Scientific, Pittsburgh, PA) pre-warmed to 34°C, for different times as mentioned in respective figures. When required, animals were recovered in 20°C incubator following heat shock, under standard condition for different time durations as described in respective figures. For experiments involving F1 progeny, eggs from control or heat-shocked mothers (P0) were collected on fresh plates and they were allowed to grow as day-one adults at 20°C incubator.

## Bleach Hatching

*C. elegans* populations containing 250-300 gravid adults were generated by passaging L4s, as described above, and waiting for five days till the majority of the animals were gravid day-one adults. Animals were washed off the plates with 1X PBS and pelleted by centrifuging at 5000 rpm for 30 s. The PBS was removed carefully, and worms were gently vortexed in presence of bleaching solution (250 µl 1N NaOH, 200 µl standard bleach, 550 µl sterile water) until all the worm bodies were dissolved (approximately 5–6 min). The eggs were pelleted by centrifugation (5000 rpm for 45 s) and bleaching solution was removed. Eggs were washed with sterile water 3-4 times and then counted. Care was taken to ensure that all the embryos hatched following this treatment. The eggs were seeded on fresh OP50 or RNAi plates (∼200 eggs/plate for chromatin immunoprecipitation) and allowed to grow as day-one-adults under standard condition (20°C).

## Egg Count

Wild-type (N2) L4s were picked on fresh OP50 plates the day before the experiment. After 24-26hr, one-day-old animals were either heat shocked at 34°C for 5, 30 or 60 min in the water bath as described above or left untreated (control). Heat-shocked animals were recovered at 20°C following heat shock and 5 animals were moved to new OP50 plates at different time intervals (0hr, 2hr, 4hr and 8 hr.) and eggs were collected for different time intervals. As described earlier, eggs are extremely susceptible to heat stress and >50% eggs laid between 0-2 hr. post heat shock didn’t hatch (*1*). Finally, eggs laid between 2-4hr, 4-8hr and 8-12hr post heat shock were counted. Eggs laid by control animals for the same duration were also collected and counted.

## Thermotolerance assay

Control and heat-shocked (34°C for different time durations as mentioned in respective figures and figure legends) mothers of different genotypes were allowed to lay eggs for different time intervals (2–4 hr. or 4—8 hr. or 8—12 hr. post-heat shock as described in respective figures and figure legends) and then all P0 mothers were removed from the plates. Plates were then kept in an incubator at 20°C until the F1 progeny become day-one adults and the progeny were then subjected to a prolonged (2 hr. 15 min) heat exposure of 37°C and then allowed to recover at 20°C overnight. This condition was chosen after prior experiments to titrate death of control, non-heat shocked progeny to <∼50% to prevent ceiling effects. The percent animals that survived the prolonged heat shock was scored 24 hrs. later.

## RNA interference

RNAi experiments were conducted using the standard feeding RNAi method. Bacterial clones expressing the control (empty vector pL4440) construct and the dsRNA targeting different *C. elegans* genes were obtained from the Ahringer RNAi library (*2*) now available through Source Bioscience (https://www.sourcebioscience.com/errors?aspxerrorpath=/products/life-science-research/clones/rnai-resources/c-elegans-rnai-collection-ahringer/). All RNAi clones used in experiments were sequenced for verification before use. For RNAi experiments, RNAi bacteria with empty (pL4440 vector as control) or RNAi constructs were grown overnight in LB liquid culture containing ampicillin (100 µg/ml) and tetracycline (12.5 µg/ml) and then induced with IPTG (1 mM) for 2 hr. before seeding the bacteria on NGM plates supplemented with ampicillin (100 µg/ml), tetracycline (12.5 µg/ml) and IPTG (1 mM). Bacterial lawns were allowed to grow for 48 hrs. before the start of the experiment. RNAi-induced knockdown was conducted by (a) dispersing the bleached eggs onto RNAi plates or (b) feeding L4 animals for 24 hrs. (as they matured from L4s to 1-day-old adults) or (c) feeding animals for over one generation, where second-generation animals were born and raised on RNAi bacterial lawns (for *hsf-1* RNAi). *mkcSi13 II; rde-1(mkc36) V* (DCL569) worms were used for germline-specific RNAi experiments (*3*). For experiments with F1 progeny, mothers (P0) were grown on RNAi plates and then eggs were collected on plates containing OP50 bacteria, so, worms in F1 generation were grown on regular plate until they were day-one-adults. Unless otherwise mentioned, day-one adults were used for experiments.

## Scoring of polyglutamine (polyQ) aggregates

Transgenic animals (*rmIs132* [*unc-54*p::Q35::YFP] expressing polyglutamine (polyQ) expansion proteins fused with fluorescent proteins in their muscle were used for the experiment. Animals growing on OP50 bacteria or RNAi bacteria (with empty L4440 vector or *met-2* RNAi construct) were either left untreated (control) or heat-shocked at 34°C for indicated time duration. Eggs were collected from control and heat-shocked P0 mothers at different time intervals: (a) eggs laid between 2-4hr post 5min heat shock, (b) eggs laid between 4-8hr post 30min heat shock and (c) eggs laid between 8-12hr post 60min heat shock. These eggs were allowed to grow till day-one adults (F1 progeny) under standard condition (20°C). Aggregates were scored in one-day-old adult and then every 24h interval over a period of 4 days using a Leica fluorescent stereomicroscope (MZFLIII) with the EYFP filter set (excitation, 510/20; emission, 560/40). Aggregates were recognized visually, based on experience from FRAP regarding which foci did not recover following photobleaching. Aggregates too close to be resolved under the stereomicroscope, or smaller satellite aggregates scattered near a large aggregate were accounted as one. This system was consistently maintained for all experiments. The number of aggregates in each worm was scored blind, by independent investigators and yielded very similar results. To quantify Q35 aggregation in the body wall muscle cells, all visible aggregates in approximately 100 animals was scored, corresponding to more than three biological samples.

## Immunostaining of dissected worms

Immunostaining of dissected worms was performed as previously described (*1*). Control or heat-shocked (34°C for indicated time) animals were transferred in 1X PBS (pH 7.4) on a cover glass (catalog no. 3800105, Leica biosystem). Worms were quickly dissected with a blade (catalog no. 4-311, Integra Miltex), and fixed with freshly prepared 4% paraformaldehyde (catalog no. 15710, Electron Microscopy Science) for 6 min. A charged slide (Superfrost Plus, catalog no. 12-550-15, Thermo Fisher Scientific) was then placed over the cover glass, and immediately placed on a pre-chilled freezing block on dry ice for at least 10 min. The cover glass was quickly removed, and the samples were post-fixed with pre-chilled 100% methanol (−20°C) for 2 min. The slides were washed in 1X PBST (1X PBS with 0.1% Tween-20), and then blocked in 1X PBST with 0.5% BSA for 1 hr. Samples were incubated at 4°C overnight with primary antibody diluted in 1X PBST. The next day, slides were rinsed in 1X PBST, and incubated with secondary antibody for 2 hr. After washing samples in 1x PBST, they were mounted in Vectashield mounting medium with DAPI (catalog no. H-1200, Vector Laboratories). Dilution factors for antibodies are: mouse anti-FLAG M2 antibody (catalog no. F-3165, Sigma Aldrich)-1:2000; Rabbit anti-HA antibody (catalog no. ab9110, Abcam)-1:500; Rabbit anti-H3K9me2 antibody (catalog no. C15410060, Diagenode)-1:500; Alexa Fluor 647 goat anti-mouse IgG (H+L) antibody (catalog no. A-21236, Thermo Fisher Scientific)-1:500; Alexa Fluor 488 goat anti-rabbit IgG (H+L) antibody (catalog no. A-11008, Thermo Fisher Scientific)-1:500. Imaging of slides was performed using a Leica TCS SPE Confocal Microscope (Leica) with a 63X oil objective. LAS AF software (Leica) was used to obtain and process Z-stack. The LAS AF software was also used to quantify the number of oocytes labelled by H3K9me2 (Threshold setting was adjusted to yield a similar low reading for the control of each experiment). For analysis of colocalization between HSF-1 and MET-2, Image J software was used; a line was drawn on one-section of Z-stack, and then intensities along the line for each channel were measured.

## Immunostaining of Embryos

Embryos were obtained from control and heat-shocked (34°C for 5 and 60min followed by 5 hrs. recovery at 20°C) animals. Worms were dissected in 1X PBS (pH7.4) on a cover glass, and a charged slide was placed over the cover glass. Samples were immediately frozen on pre-chilled freezing block in liquid nitrogen for at least 10 min, and then immersed in liquid nitrogen for 2 min. After freezing for an additional 1 hr. at −80°C, the cover glass was quickly removed. Embryos were subsequently fixed at −20°C in pre-chilled 100% methanol and 100% acetone for 30 min each. The slides were then incubated successively in pre-chilled 70%, 50%, and 30% ethanol (−20°C) for 1 min each and washed twice in 1X PBST. The remaining process from blocking to imaging was performed as described for imaging of dissected worms. Rabbit anti-H3K9me2 antibody (catalog no. C15410060, Diagenode)-1:500 and Alexa Fluor 488 goat anti-rabbit IgG (H+L) antibody (catalog no. A-11008, Thermo Fisher Scientific)-1:500 were used. All images were quantified using Image J; H3K9me2 intensity for each nucleus was measured as raw integrated density divided by area.

## DAF-16::GFP nuclear localization

Transgenic animals expressing DAF-16::GFP (*zIs356*) were used for the experiment. Worms (P0) were either left untreated (control) or heat-shocked at 34°C for 5 and 60 min. Eggs from control and heat shocked worms (2-4 hr. eggs post 5min heat shock and 8-12 hr. eggs post 60min heat shock) were collected on fresh NGM plates seeded with OP50 bacteria and allowed to grow at 20°C as day-1 adults. Day-one progeny (F1) were mounted on a 2% agar pad containing a drop of 10mM levamisole. All worms were imaged within 10 min using a Leica TCS SPE Confocal Microscope (Leica) with a 63X oil objective. LAS AF software (Leica) was used to obtain one section of image as well as score the number of worms that show DAF-16 nuclear localization.

## RNA extraction and quantitative real-time PCR (qRT-PCR)

Unless otherwise mentioned, RNA was extracted from day-one adults. Animals were either passaged as L4s, 24 hrs. before harvesting (for P0 animals) or harvested as day-one adults (for F1 progeny) at densities of 20-30 worms/plate. RNA was extracted as described earlier (*4*). Briefly, RNA samples were harvested in 50 µl of Trizol (catalog no. 400753, Life Technologies) and snap-frozen immediately in liquid nitrogen. Samples were thawed on ice and 200 µl of Trizol was added, followed by brief vortexing at room temperature. Samples were then lysed using a Precellys 24 homogenizer (Bertin Corp.) to lyse the worms completely. RNA was then purified as detailed with appropriate volumes of reagents modified to 250 µl of Trizol. The RNA pellet was dissolved in 17 µl of RNase-free water. The purified RNA was then treated with deoxyribonuclease using the TURBO DNA-free kit (catalog no. AM1907, Life Technologies) as per the manufacturer’s protocol. cDNA was generated by using the iScript cDNA Synthesis Kit (catalog no. 170–8891, Bio-Rad). qRT-PCR was performed using PowerUp SYBR Green Master Mix (catalog no. A25742, Thermo Fisher Scientific) in QuantStudio 3 Real-Time PCR System (Thermo Fisher Scientific) at a 10 µl sample volume, in a 96-well plate (catalog no. 4346907, Thermo Fisher Scientific). The relative amounts of mRNA were determined using the ΔΔ*C*_t_ method for quantitation. We selected *pmp-3* as an appropriate internal control for gene expression analysis in *C. elegans*.

All relative changes of mRNA were normalized to either that of the wild-type control or the control for each genotype (specified in figure legends). Each experiment was repeated a minimum of three times. For qPCR reactions, the amplification of a single product with no primer dimers was confirmed by melt-curve analysis performed at the end of the reaction. Reverse transcriptase-minus controls were included to exclude any possible genomic DNA amplification. Primers were designed using Primer3 software and generated by Integrated DNA Technologies. The primers used for the qRT-PCR analysis are listed below:

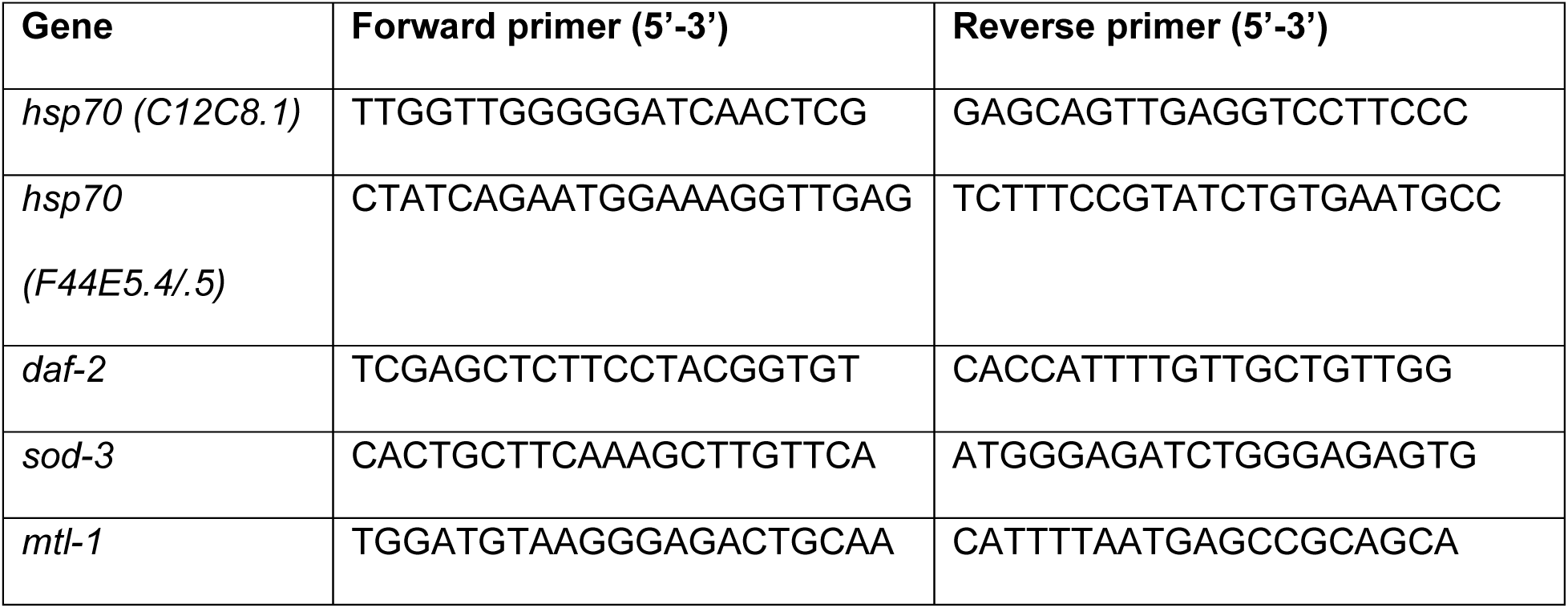
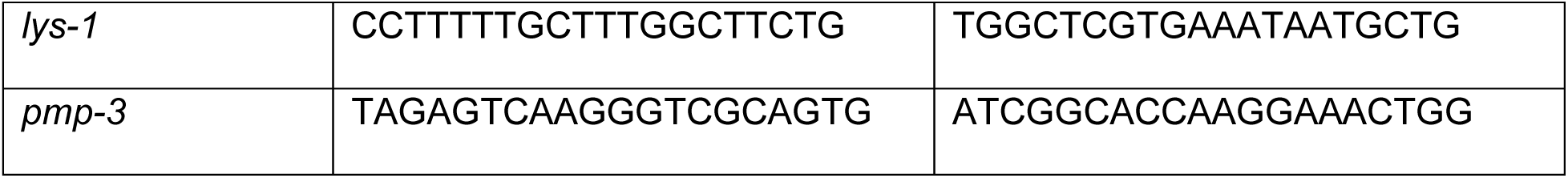

## Co-immunopreciptation (Co-IP)

∼1,000-1,200 day-1 adults per condition (control or heat shock at 34°C for 30 min) were collected in M9 buffer and snap frozen in liquid nitrogen. Worm pellets were resuspended with equal volume of pre-chilled 2x lysis buffer [60mM HEPES (pH7.4), 100 mM KCl, 4 mM MgCl, 0.1% Triton X-100, 10% glycerol, supplemented with 2 mM DTT and protease inhibitor cocktail] (*5*). After lysing with a Precellys 24 homogenizer (Bertin Corp.), samples were centrifuged at 200g for 5 min at 4°C to remove worm debris. 10% of worm lysates was snap frozen to use as input. For immunoprecipitation of HA tagged MET-2, the remaining lysates were incubated at 4°C for 2 hours with mouse anti-HA magnetic bead (catalog no. 8837, Pierce). Before incubation, the beads were pre-cleared with pre-cold 1X lysis buffer. Anti-mouse IgG antibody (catalog no. ab188776, Abcam) was used as negative control. Before incubation with samples, anti-mouse IgG antibody was conjugated with Protein A/G magnetic beads (catalog no. 88802, Pierce) at 4°C for 1 hr. Beads were then washed with pre-chilled wash buffer [30 mM HEPES, pH = 7.4, 100 mM KCl, 2mM MgCl, 0.1% Triton X-100, supplemented with 1 mM DTT], and mixed with 4X Laemmli sample buffer (catalog no. 1610737, Bio-Rad) supplemented with 10% β-mercaptoethanol. Immunoprecipitated protein mixture was eluted and denatured by boiling, and analyzed by western blotting with mouse anti-FLAG M2 antibody (1:1000; catalog no. F1804, Sigma Aldrich) and rabbit anti-HA antibody (1:1000; catalog no. ab9110, Abcam).

## Western Blotting

Western blot analysis was performed with adult day-1 animals. For protein analysis, 15–50 worms (depending on the experimental requirement) were harvested in 15 µl of 1X PBS (pH 7.4), and then 4X Laemmli sample buffer (catalog no. 1610737, Bio-Rad) supplemented with 10% β-mercaptoethanol was added to each sample. Samples were then boiled for 30 min. Whole-worm lysates were resolved on SDS-PAGE gels and transferred onto nitrocellulose membrane (catalog no. 1620115, Bio-Rad). Membranes were blocked with Odyssey Blocking Buffer (part no. 927–50000, LI-COR). Immunoblots were imaged using LI-COR Odyssey Infrared Imaging System (LI-COR Biotechnology, Lincoln, NE). Mouse anti-FLAG M2 antibody (catalog no. F1804, Sigma Aldrich) was used to detect HSF-1::FLAG and C12C1.8::FLAG. Rabbit anti-HA antibody (catalog no. ab9110, Abcam) was used to detect MET-2::HA and HSP-1::HA. Mouse anti-polyglutamine antibody (catalog no. P1874, Sigma Aldrich) was used to detect polyQ expression in Q35::YFP animals (*6*). Rabbit anti-HSP90 antibody (catalog no. 4874S) was used to detect DAF-21 level. Mouse anti-H3K9me2 antibody (catalog no. ab1220, Abcam) was used to detect dimethylated H3K9 and rabbit anti-histone H3 antibody (catalog no. ab1791, Abcam) was used to detect total histone H3 level. Mouse anti-α-tubulin primary antibody (catalog no. AA4.3), developed by C. Walsh, was obtained from the Developmental Studies Hybridoma Bank (DSHB), created by the National Institute of Child Health and Human Development (NICHD) of the National Institute of Health (NIH), and maintained at the Department of Biology, University of Iowa. The following secondary antibodies were used: Sheep anti-mouse IgG (H and L) Antibody IRDye 800CW Conjugated (catalog no. 610-631-002, Rockland Immunochemicals) and Alexa Fluor 680 goat anti-rabbit IgG (H+L) (catalog no. A21109, Thermo Fisher Scientific). Protein expression (relative to α-tubulin levels) in different samples was measured using the LI-COR Image Studio software. Fold change of protein levels was calculated relative to wild-type/untreated controls.

## Chromatin immunoprecipitation

Chromatin immunoprecipitation (ChIP) was performed as described earlier (*1*). For ChIP in P0 animals (mothers), about ∼500 day-1 animals per condition (control or heat shock at 34°C for indicated time) were obtained by washing off two plates of bleach hatched animals. For ChIP in F1 animals (progeny), ∼400-500 F1 worms per condition were collected from P0 animals (control or heat shock at 34°C for indicated time and then eggs were collected as mentioned in the figures/ figure legends). Worms were washed with 1X PBS (pH 7.4), and cross-linked with freshly prepared 2% formaldehyde (catalog no. 252549, Sigma Aldrich) at room temperature for 10 min followed by addition of 250 mM Tris (pH 7.4) at room temperature for 10 min. Samples were then washed three times in ice-cold 1X PBS supplemented with protease inhibitor cocktail and snap-frozen in liquid nitrogen. The worm pellet was resuspended in FA buffer [50 mM HEPES (pH 7.4), 150 mM NaCl, 50 mM EDTA, 1% Triton-X-100, 0.5% SDS and 0.1% sodium deoxycholate], supplemented with 1 mM DTT and protease inhibitor cocktail. The suspended worm pellet was lysed using a Precellys 24 homogenizer (Bertin Corp.), and then sonicated in a Bioruptor Pico Sonication System (catalog no. B0106001, Diagenode) (15 cycles of 30 s on/off). For all HSF-1 ChIP experiments, endogenous HSF-1 was immunoprecipitated with anti-FLAG M2 magnetic bead (catalog no. M-8823, Sigma-Aldrich). Beads were first pre-cleared with chromatin isolated from wild-type worms not having any FLAG tag and salmon sperm DNA (catalog no. 15632–011, Invitrogen). Worm lysate was incubated at 4°C overnight with the pre-cleared FLAG beads. For all other ChIP experiments, Protein A/G Magnetic Beads (catalog no. 88802, Pierce) pre-cleared with salmon sperm DNA was used. Pre-cleared lysate was incubated at 4°C overnight with anti-H3K9me2 antibody (*7*) (catalog no. ab1220, Abcam) and then pre-cleared magnetic bead was added and incubated for another 3–4 hr. Beads were washed with low salt, high salt and LiCl wash buffers and then eluted in buffer containing EDTA, SDS and sodium bicarbonate. The elute was incubated with RNase A and then de-crosslinked overnight in presence of Proteinase K. The DNA was purified by ChIP DNA purification kit (catalog no. D5205, Zymo Research). qPCR analysis of DNA was performed as described above using primer sets specific for different target genes. Promoter region of *syp-1* was amplified for all HSF1-ChIP experiments to quantify non-specific binding of HSF-1. For all ChIP experiments, 10% of total lysate was used as ‘input’ and chromatin immunoprecipitated by different antibodies were expressed as % input values. All relative changes were normalized to either that of the wild-type control or the control of each genotype (specified in figure legends) and fold changes were calculated by ΔΔ*C*_t_ method. The primers used for ChIP experiments, and the expected amplicon sizes are as follows:

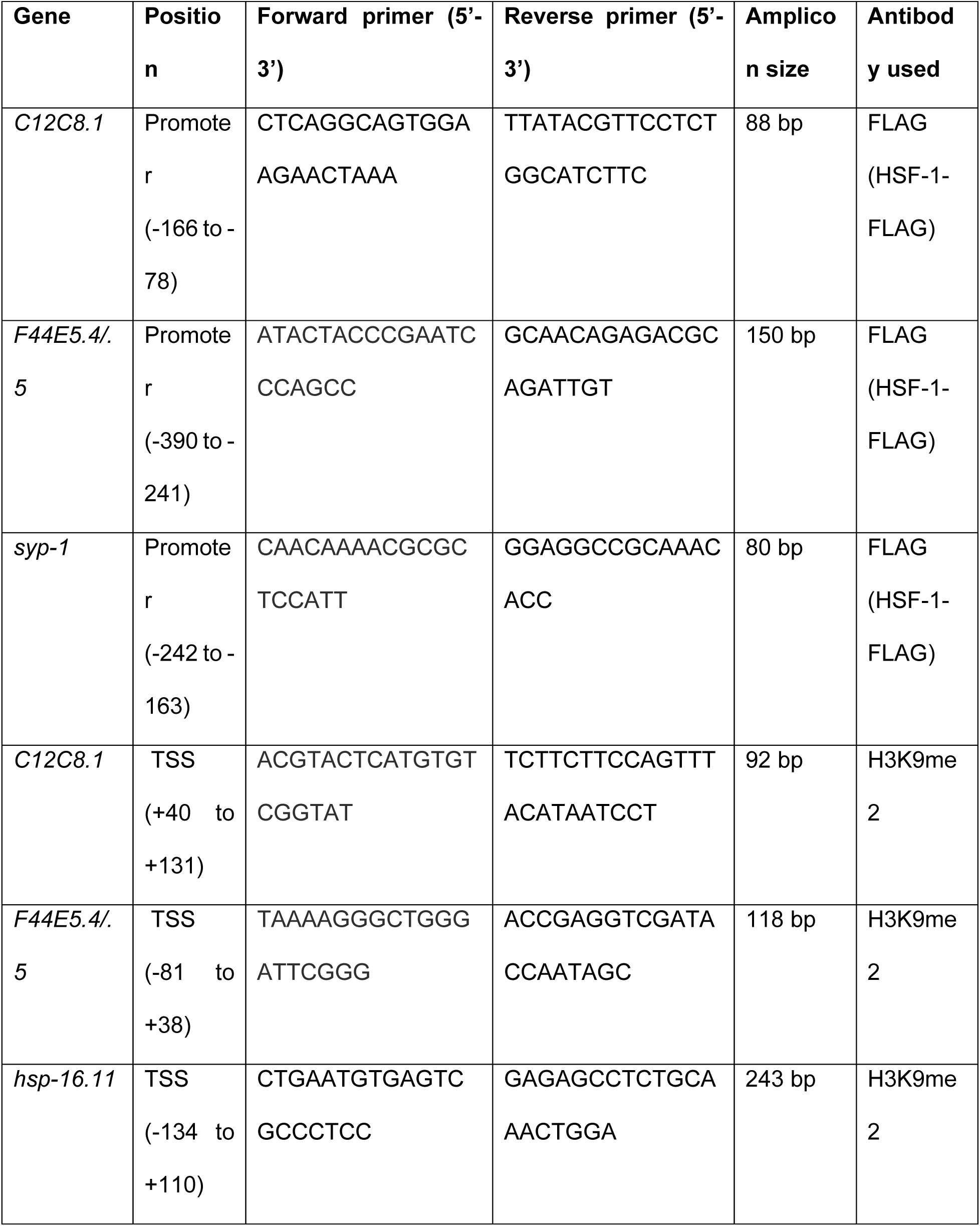
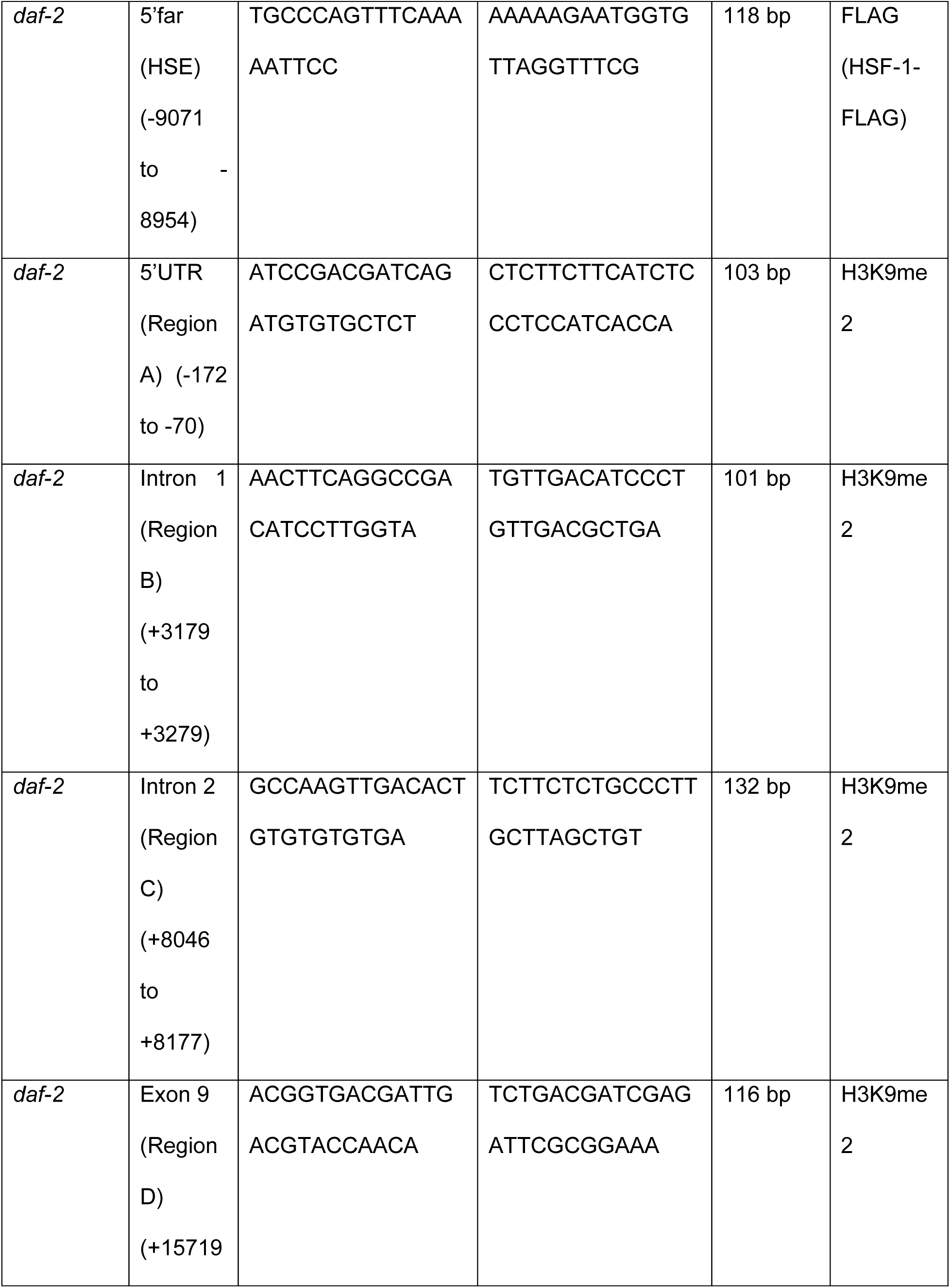
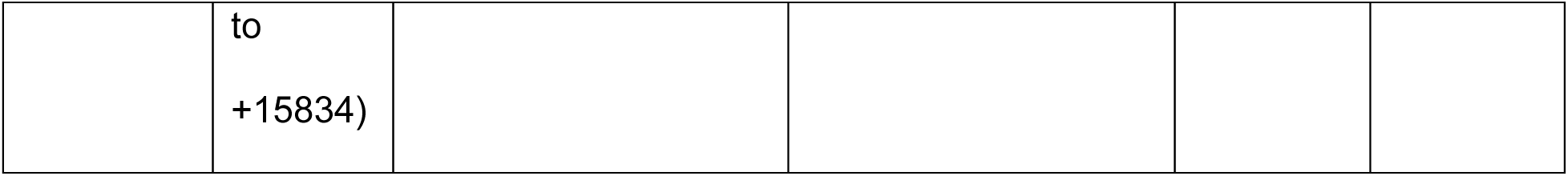

